# Developing a temperature-inducible transcriptional rheostat in *Neurospora crassa*

**DOI:** 10.1101/2022.11.24.517854

**Authors:** Cyndi Tabilo-Agurto, Verónica Del Rio-Pinilla, Valeria Eltit-Villarroel, Alejandra Goity, Felipe Muñoz-Guzmán, Luis F. Larrondo

**Affiliations:** Departamento de Genética Molecular y Microbiología, Facultad de Ciencias Biológicas, Pontificia Universidad Católica de Chile, ANID-Millennium Science Initiative-Millennium Institute for Integrative Biology (iBIO), Alameda 340. Santiago 8331150, Chile

**Author notes:** Address correspondence to Luis F. Larrondo. Present address: Felipe Muñoz-Guzmán, Departamento de Biología, Facultad de Química y Biología, Universidad de Santiago de Chile, Santiago, 9170022, Chile. Cyndi Tabilo-Agurto and Verónica Del Rio-Pinilla contributed equally to this work. Author order was determined based on the date they started their participation in this project.

**Keywords:** *hsp* promoters, heat shock, synthetic promoter, inducible promoter, *Neurospora crassa*

## Abstract

Heat shock protein (*hsp*) encoding genes, part of the highly conserved Heat Shock Response (HSR), are known to be induced by thermal stress in several organisms. In *Neurospora crassa*, three *hsp* genes, *hsp30, hsp70*, and *hsp80*, have been characterized; however, the role of defined *cis*-elements in their response to discrete changes in temperature remains largely unexplored. To fill this gap, while also aiming to obtain a reliable fungal heat-shock inducible system, we analyzed different sections of each *hsp* promoter, by assessing the expression of real-time transcriptional reporters. Whereas all three promoters, and their resected versions, were acutely induced by high temperatures, only *hsp30* displayed a broad range of expression and high tunability amply exciding other inducible promoter systems existing in Neurospora, such as Quinic acid- or light-inducible ones. As proof of concept, we employed one of these promoters to control the expression of *clr-2*, which encodes for the master regulator of Neurospora cellulolytic capabilities. The resulting strain fails to grow on cellulose at 25°C, whereas it robustly grows if heat shock pulses are delivered daily. Additionally, we designed two *hsp30* synthetic promoters and characterized these, as well as the native promoters, to a gradient of high temperatures, yielding a wide range of responses to thermal stimuli. Thus, Neurospora *hsp30*-based promoters represent a new set of modular elements that can be used as a transcriptional rheostat to adjust the expression of a gene of interest or for the implementation of regulated circuitries for synthetic biology and biotechnological strategies.

**Importance:** Timely and dynamic response to strong temperature rises is paramount for organismal biology. At the same time, inducible promoters are a powerful tool for fungal biotechnological and synthetic biology endeavors. In this work, we analyzed the activity of several *N. crassa* heat shock protein (*hsp*) promoters upon a wide range of temperatures, observing that *hsp30* exhibits remarkable sensitivity and dynamic range of expression as we chartered the response of this promoter to subtle increases in temperature, while also building synthetic promoters based on *hsp30 cis*-elements. As proof of concept, we analyzed the ability of *hsp30* to provide tight control of a central process such as cellulose degradation. While this study provides an unprecedented description of the regulation of the *N. crassa hsp* genes it also contributes with a noteworthy addition to the molecular toolset of transcriptional controllers in filamentous fungi.

## Introduction

The filamentous fungus *Neurospora crassa* has been used as a model organism for the molecular dissection of diverse complex biological processes, such as cellulose degradation [1,2], gene silencing [3–5], circadian rhythms [6–9], and photobiology [10]. The broad set of available molecular tools in this organism is also extensive, including a knockout collection [11], selectable markers [12–15], CRISPR/Cas9 technologies [16], and inducible/constitutive promoters [10,17,18].

Different promoters induced by chemical signals have been developed in *N. crassa*, such as ones that can respond to glucose [19], Quinic acid [20,21], nitrogen [22–24], and copper [24,25]. Nevertheless, the utilization of physical cues as inducing signals has mainly focused on the use of promoters responding to light [18,26,27], whereas other signals, like temperature, have seldom been employed as modulators of transcriptional units in this organism.

The Heat Shock Response (HSR), is an evolutionary conserved protective mechanism triggered by high temperatures, which can also be induced by other stresses [28]. Inside cells, the HSR leads to, among other cellular changes, an intense and rapid synthesis of proteins acting as chaperons called Heat Shock Proteins (HSPs). These proteins are well conserved in terms of features and functions, and some of them have been also shown to play roles in normal development [28,29]. In eukaryotes, the *hsp* promoters have revealed the presence of consensus heat-responsive sequences known as heat shock elements (*hse*; 5’-nTTCnnGAAnnTTCn-3’). Heat Shock Factors (HSF) are the proteins in charge of recognizing *hse* boxes, exhibiting similar DNA binding domains across eukaryotes [30].

The *hsp* promoters and their expression profiles have been well studied in diverse model organisms, including plants [31,31], mammals [33,34], and fungi [35–38]. In several cases, *hsp* promoters have been successfully utilized for spatial and temporal control of gene expression [39–41], to promote heat stress tolerance in distinct organisms [42–45], in heterologous protein and chemical production [46,47] or even utilized for synthetic circuits-based biosensors [48].

In *N. crassa*, three genes encoding for the major HSP from each family have been described: *hsp30* (NCU09364) [49], *hsp70* (NCU09602) [50], and *hsp80* (NCU04142) [51]. It has been reported that these three *hsp* genes are expressed in response to high temperatures [50,52,53], and that they bear putative *hse* regulatory elements in their promoter regions [49-51,54]. Despite this, further characterization of the *hsp* promoters in *Neurospora* has not been systematically conducted, nor minimal aspects such as dynamic ranges of expression and their tunability by discrete temperature changes have been studied. Such analyses are not only relevant to better understand how *Neurospora* responds to thermal stimuli but can also yield valuable information on which *hsp* promoter(s) can be adopted as viable and versatile inducible systems.

In this work, we sought to characterize the transcriptional response of the *hsp30, hsp70*, and *hsp80* promoters utilizing a destabilized codon-optimized luciferase, a well-known reporter for transcriptional dynamics in *Neurospora crassa* [55,56]. Indeed, the addition of a degron (PEST sequence) to firefly luciferase turns this real time reporter into a great system to dissect promoters of interest, including their range of inducibility upon cognate stimuli. Thus, we assessed the regulation conferred by the full and resected version of each promoter upon exposure to different temperatures. Because of their highly tunable regulation and low basal level of expression, we selected *hsp30*-derived ones to further delve into their expression dynamics by exposing them to a gradient of high temperatures and a variety of treatment times. The end result is an accurate profile of their responses to diverse temperature stimuli. In addition, and as a proof of concept of their applicability, we utilized a resected *hsp30* promoter to control the expression of *clr-2*, which encodes the master transcription factor involved in cellulose degradation, resulting in heat-shock conditional growth. Finally, we designed two synthetic promoters based on multiple *hsp30* (*SP30*) putative heat response elements in order to generate modular versions of these sequences to avoid RIP and also to facilitate future synthetic biology strategies.

*In toto*, the results provide new and detailed data on *hsp* responses to temperature in *Neurospora*, while also establishing *hsp30*-derived systems as versatile transcriptional rheostats for graded gene expression, expanding the existing repertoire of inducible promoters in filamentous fungi.

## Results

### Functional analysis of hsp promoters

In previous reports, the sequences of the *hsp30, hsp70*, and *hsp80* promoters have been succinctly described [49–51]. For *hsp30* and *hsp70*, four and two putative *hse* have been proposed (Figure 1a, 1b), albeit none of them has a perfect match with the described consensus sequence (Table S2, S3). For *hsp80*, previous studies did not report sequences resembling the consensus *hse* and, instead, described temperature response elements (*tre*) (Figure 1c), which the authors proposed might allow the expression of this gene upon heat shock [51]. However, the functionality of none of the regions containing these putative *cis*-elements, in any of these promoters, has been experimentally confirmed.

**Figure 1.**
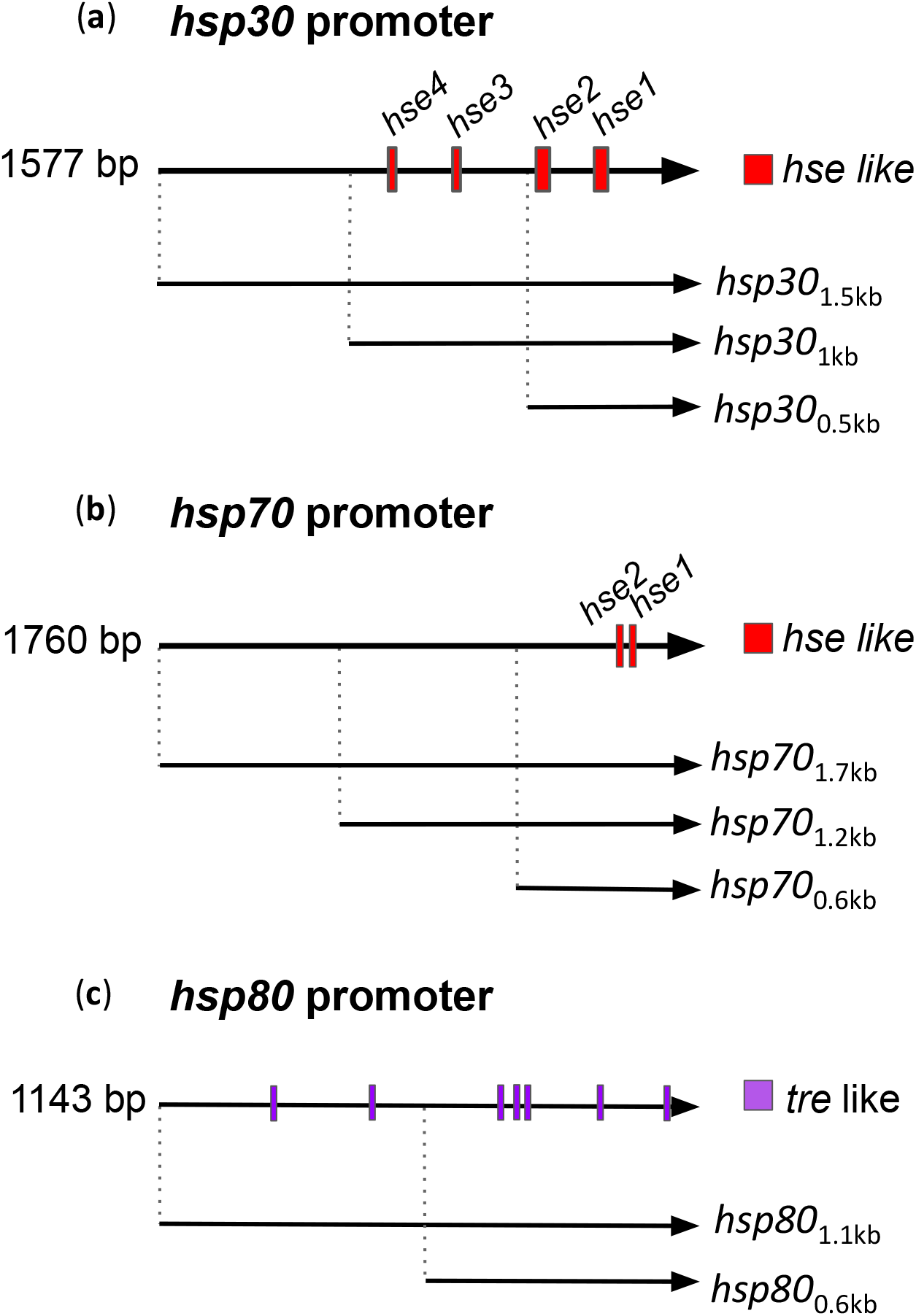
Putative transcriptional heat shock regulatory elements present in the *hsp* promoters. (**a** to **c**) Scheme of *hsp30, hsp70*, and *hsp80* promoters where the putative transcriptional regulatory elements are indicated [49–51,54]. The *hsp30* and *hsp70* promoters contain putative heat shock elements (*hse*, red boxes), while the *hsp80* promoter bears putative temperature responsive elements (*tre*, purple boxes). We analyzed the indicated section, upstream from the ORF (arrow head). The dimension of the boxes and lines represent the size of transcriptional regulatory elements and the promoter region, respectively.

To advance such functional analyses, we generated an array of reporter strains spanning different promoter dissections of the abovementioned *hsp* genes, controlling the expression of a destabilized firefly luciferase. The dissected regions consider different lengths of upstream sequence (relative to the ORF), that were selected depending on the presence of the putative heat-responsive elements (Figure 1). Thus, we generated 8 dissections: three from full-length promoters for each *hsp* (*hsp30*_1.5kb_, *hsp70*_1.7kb_, *hsp80*_1.1kb_) based on the previously described sequences [49-51,54], plus two resected versions for *hsp30* (*hsp30*_1kb_ and *hsp30*_0.5kb_), two for *hsp70* (*hsp70*_1.2kb_ and *hsp70*_0.6kb_), and one resected section for *hsp80* (*hsp80*_0.6kb_). Both the full-length and the 1 kb section of *hsp30* share the same four putative *hse*, while the smallest region has only two of those boxes. The *hsp70* promoter region contains the two previously proposed *hse* (Figure 1b), whereas the *hsp80*_0.6kb_ section keeps five out of the seven potential *tre*-like elements (Figure 1).

As an exploratory analysis, we grew the reporter strains in constant light conditions (LL) for 24 hours and then, using a CCD camera, recorded the luciferase activity of each promoter in darkness (DD) at 25°C (Figure 2). After 48 hours, we exposed them to 1-hour heat shock at three different temperatures: two high ones (35°C and 45°C) and one closer to *Neurospora* laboratory growth conditions (30°C). We observed that the three full-promoters had similar fast and strong induction at 45°C, displaying only a reduced response at 35°C, while no change was seen at 30°C (Figure 2). The resected *hsp30*_1kb_ version had a similar transcriptional profile compared to *hsp30*_1.5kb_ at all temperatures, whereas *hsp30*_0.5kb_ showed a reduced expression at 45°C, with less than half the levels of the full-promoter, and no obvious induction at 35°C (Figure 2). The shorter versions of the *hsp70* constructs showed, at all tested temperatures, a similar transcriptional profile to the full version (Figure 2). The *hsp80*_0.6kb_ region displayed equally diminished responses at both high temperatures, being about 9-times weaker than what was observed for *hsp80*_1.1kb_ at 45°C (Figure 2).

**Figure 2.**
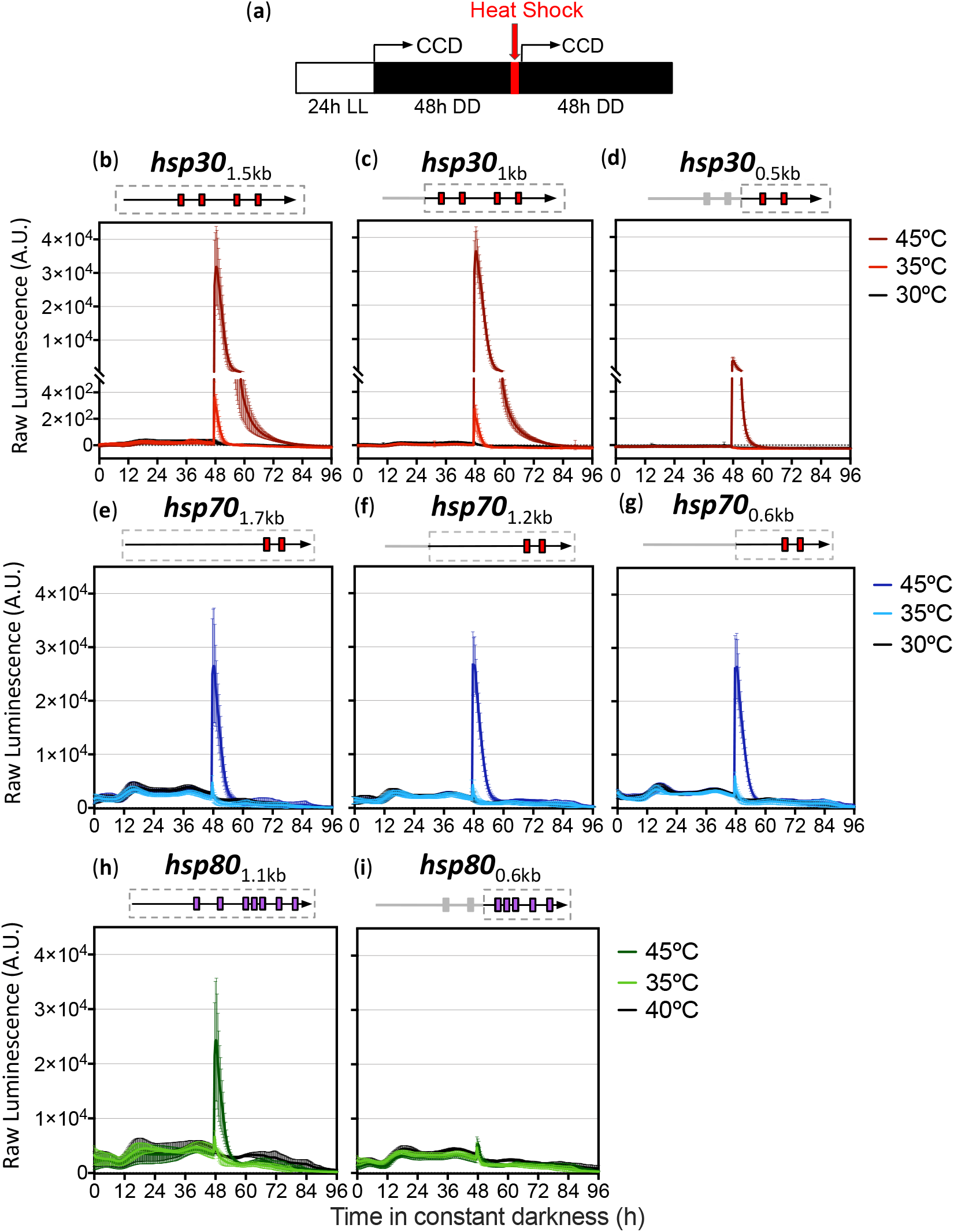
Luciferase activity profiles conferred by *hsp* promoters and resected sections upon heat shock treatment. (**a**) Description of the experimental setup. The broken arrows represent the start of the bioluminescence measurements in the CCD camera. The strains grew for 24 h at 25°C in constant light conditions (LL), and then we measured the luminescence at 25°C in constant darkness (DD). The heat-shock treatments (30°C, 35°C, and 45°C) were delivered for one hour using an incubator, after which luminescence was monitored for additional 48 h. (**b** to **i**) Luminescence levels are shown in arbitrary units (A.U.). Boxes in gray dotted lines above each chart represent the area of the promoter region being analyzed, whereas the red and purple boxes represent the putative *hse* and *tre*, respectively. Each curve corresponds to the average of four to six biological clones with three independent wells each ± standard deviation (SD) and represent the behavior in two independent experiments.

Comparing the overall expression profiles between the three *hsp* promoters, we observed that *hsp70* and *hsp80* exhibited higher basal activity at 25°C (Figure S1). In contrast, the *hsp30* promoters have basal luminescence levels that are 10 and 15 times lower, compared to the two other *hsp*, respectively, and are strongly induced after exposure to high temperatures. Indeed, *hsp30* basal levels of expression are comparable to what is obtained when examining a lowly expressed gene such as the clock gene *frequency* [55,56]. This causes the *hsp70*- and *hsp80*- based promoters to have a limited induction profile after a heat shock, measured as the fold-induction between basal and peaks levels of luciferase activity while, on the contrary, *hsp30*-based promoters (full and resected constructs) display high fold-induction ratios (Figure 3). Thus, the *hsp30*_1.5kb_ and *hsp30*_1kb_ promoters exhibit ∼10- and over 1000-fold induction after being treated at 35°C and 45°C, respectively; whereas although the *hsp30*_0.5kb_ promoter did not display a clear response at 35°C, it yielded an activation of over 100-fold when stimulated at 45°C.

**Figure 3.**
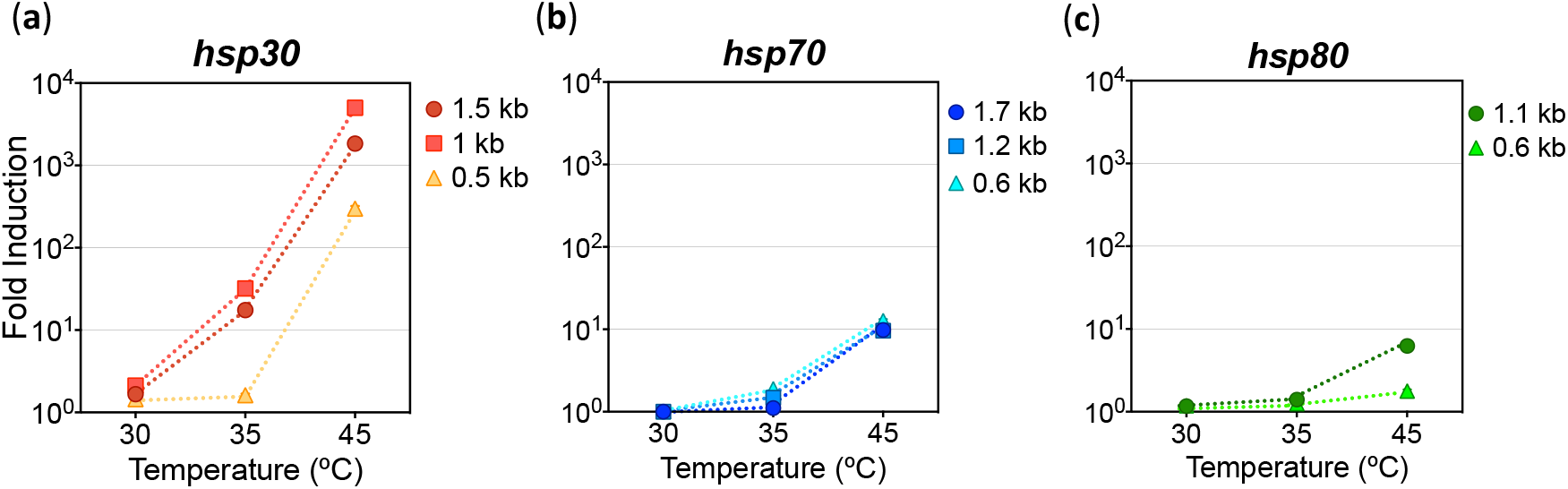
Fold induction achieved by the *hsp* promoter regions after heat shock treatments. (**a** to **c**) The fold induction (fold-change) was calculated with the maximum luciferase expression, based on the average of the three highest consecutive values respect to the background values before heat shock treatment of each promoter region. The data was obtained from Figure 2b-2i. Average fold inductions are shown.

We then sought to compare the response of *hsp30* to other well-known inducible systems normally utilized in *N. crassa* (Figure S3), such as the *qa-2* promoter, which reacts to increasing concentrations of Quinic acid (QA) [21] and the *vvd* promoter, which is activated by light [18]. In our hands *hsp30* gave inductions 10- to 1000-times higher compared to *qa-2* and *vvd* reporters, respectively (Figure S3). Importantly, we could observe that *hsp30*_1.5kb_ displayed lower background levels, and maximum response compared to the other promoters (Table S4). Additionally, regarding to the time spanned to reach the highest response, the post-versus pre-stimuli levels, or the rate of signal decay, *hsp30*_1.5kb_ showed several properties similar to the *vvd* promoter (Table S4). *hsp30*_1.5kb_ regained basal levels after stimuli ∼2-3 h longer than what was exhibited by the *vvd* promoter after a discrete light-pulse. For the *qa-2* reporter these aspects could not be evaluated as QA remains in the media after addition and, therefore, the response does not decrease after the stimulus is initiated. Importantly, the reporters gave different levels of induction in assays conducted in PCR tubes versus larger volume tubes, yet, in all cases *hsp* exceeded the *qa-2* and *vvd*-based systems. Considering all these characteristics, and the inducibility and tunability of the response of the *hsp30*-derived promoters, we further evaluated their behavior and tested their functionality.

### Detailed charting of the hsp30 promoter response to different heat shock stimuli

We then performed a detailed functional characterization of the transcriptional responses of the different *hsp30* promoters, upon discrete temperature changes within a 35-45°C range during different exposure times. To analyze this, we adopted a simple yet practical strategy that allowed us to expose the *Neurospora* reporter strains to a heat shock gradient using a 96-well plate format (Figure S2). We used arrays made by PCR tubes and a gradient-thermocycler to expose the cultures simultaneously to six different temperatures: 34°C, 36°C, 38°C, 40°C, 42°C, and 44°C; and to different treatment times: 60, 30, 15, 5 min, and 1 min. This approach allowed us to obtain faster and more accurate results than with the previous strategy (Figure 2), providing precise temperature treatment in each well.

The *hsp30* promoters under study show that the degree of the response is augmented as temperature is progressively increased, and as the duration of the stimuli is lengthened (Figure 4), behaving as transcriptional rheostats tuning responses upon changes in the strength of the stimuli (Figure 4, Figure S4). Thus, the reporters yielded increasing induction levels reaching a maximum at 15 min (Figure 4a, Figure S4), while longer heat pulses (30 and 60 min) still yielded strong responses which, in general, were lower that the peak level (15 min). Notably, a short heat pulse (1 minute) was already able to elicit robust responses, although only at the highest temperatures. Thus, the *hsp30* promoter is capable of a highly tunable response even upon short pulses of heat (Figure 4).

**Figure 4.**
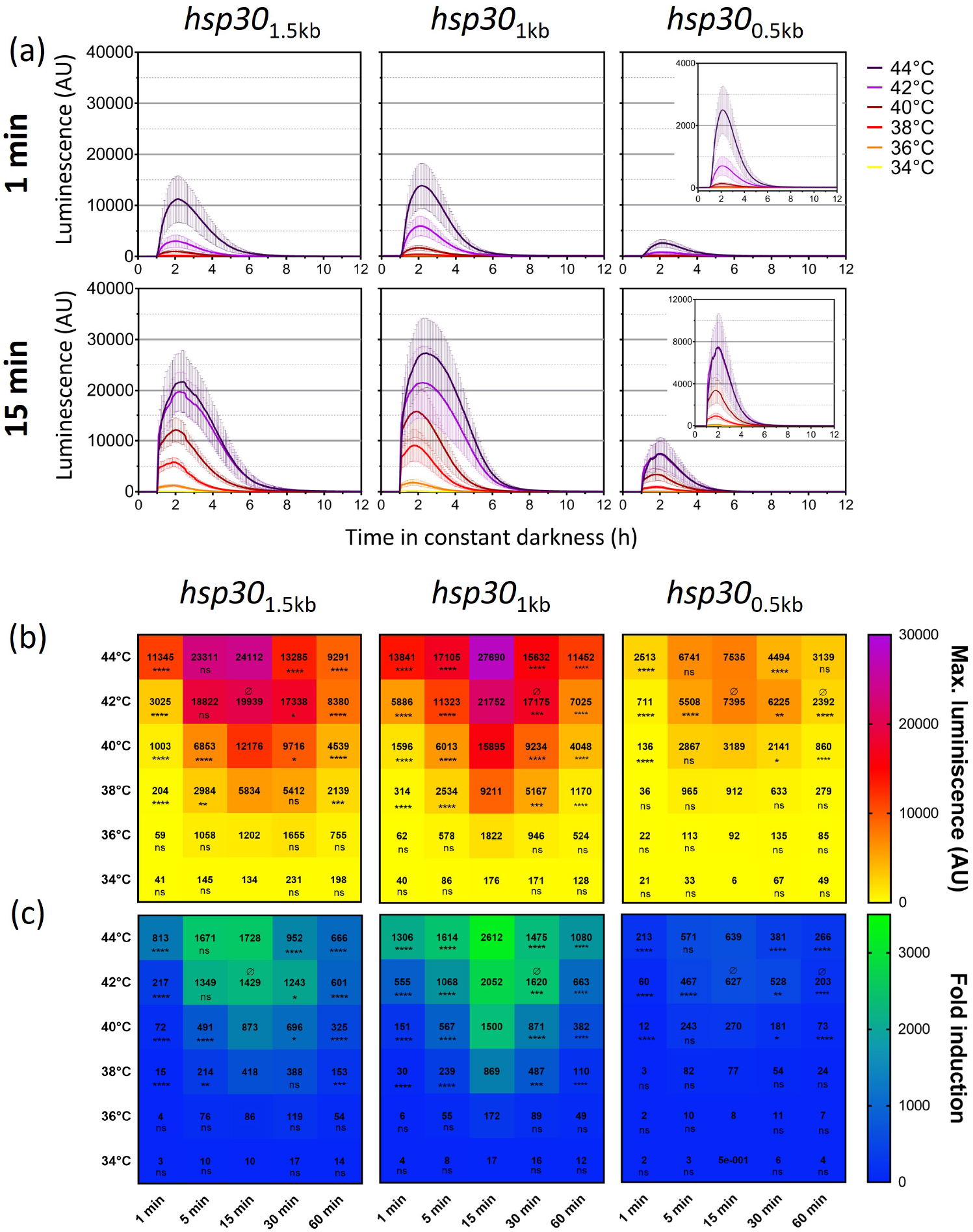
Transcriptional responses conferred by the full or resected *hsp30* promoters to a temperature gradient and different exposure times. (**a**) Activity profile of each *hsp30* promoter region to different temperatures after 1 and 15 minutes of treatment. A close-up of the *hsp30*_0.5kb_ graph is displayed on the right side. Each curve corresponds to the average of two biological clones with eight independent wells each ± standard deviation (SD) and represents the behavior in two independent experiments. (**b**) Maximum luminescence and (**c**) Fold-change obtained after all the heat-shock treatments for each *hsp30* promoter region. The maximum luminescence was defined as the average of the highest values. The fold induction was calculated with the maximum luciferase expression (shown in panel b) with respect to the average of the background values before heat shock treatment of each promoter region. The data was obtained from the luciferase activity profile shown in Figure S4. Statistical significance was performed using a two-way ANOVA plus Dunnett’s test (for time treatments all values were compared to 15 minutes: *= p<0.05; **= p < 0.01; *** = p < 0.001; **** = p < 0.0001; ns = non-significant.; and for temperature treatment all values were compared to 44°C: Ø indicates non-significant. Time significance is indicated below each value and for temperature only non-significant values are above.

The full-length promoter (*hsp30*_1.5kb_) and the *hsp30*_1kb_ section exhibited stronger inductions, compared to the *hsp30*_0.5kb_ section, reaching the maximal levels at the highest temperatures (Figure 4), confirming the trend seen in previous experiments (Figure 2). Notably, the analyses revealed that the first two promoters displayed strong activation starting at 38°C, while the smallest promoter section yields strong transcriptional responses only starting at 40°C (Figure 4, S4). As observed in the transcriptional profiles, these *hsp30* promoters generate a graded response as temperature and treatment length variables are combined, further supporting the notion of their use as transcriptional rheostats to tune the expression of genes of interest.

To better compare the abovementioned data, we calculated the fold induction achieved for each promoter under different treatment conditions (Figure 4c). This analysis confirmed that the attained induction tends to increase with higher temperatures and longer exposure times up to 15 min (Figure 4c), and that when the reporter strains are exposed for 30 or 60 min, the response is still high although less than peak levels. Thus, although we initially expected to have the largest induction at the highest temperature and longest times, this was achieved instead at 15 min. These results suggest that due to the high efficiency of thermal exchange achieved in this assay (utilizing a thermocycler), even lower applied temperature and a shorter exposure time suffice to generate a maximal response, whereas higher temperatures (for prolonged times), may lead to a detrimental effect on cellular function. Despite the different number of putative regulatory elements between the longest and the shortest promoter regions, each of the tested sections displayed a wide range of activities, confirming accurate and progressive responses to the intensity and duration of the thermal stimulus.

### Temperature-conditioned control of a N. crassa catabolic proccess

With the characterization of the dynamic regulation of the *hsp30*-derived promoters, we sought to test one of them in its ability to control a gene of interest, of relevance in fungal physiology. The *hsp30*_1kb_ region showed a similar response pattern to the full-length promoter and, while these promoters offer a great inducible system, working with long DNA segments in *Neurospora* can trigger ***r***epeated ***i***nduced ***<u>p</u>***oint (RIP) mutation during a sexual cross, which can lead to alteration of the endogenous as well as the additional copy of DNA [57]. To minimize such problems, we decided to use the *hsp30*_0.5kb_ section, which is closer to the ∼400 bp limit at which RIP starts occurring. While choosing this smaller promoter compromises levels of expression, it still provides a good dynamic range of regulation. Also, having a shorter promoter region makes it easier to use it in combination with other transcriptional modules.

*N. crassa* is known to possess great plant cell wall decomposition capabilities [58,59], having more than 400 proteins with Carbohydrate-active enzymes domains [60]. Several of the genes involved in the underlying regulatory network, as well as the master controllers of the system, have been identified [2,61]. Hence, *Neurospora* is a great model organism to dissect key aspects of industrial production of second-generation bioethanol [62–65]. One relevant strategy to control cellulase production is modulating the expression of one of the key genes in this pathway: the one encoding for the transcription factor CLR-2 (NCU08042) [66]. Thus, we aimed to command, through high-temperature, the expression of *clr-2* by putting it under the control of the *hsp30*_0.5kb_ promoter.

We evaluated the capability of the engineered strains to grow in media with Avicel (crystalline cellulose) as the sole carbon source, knowing that *clr-2* expression is needed to induce cellulolytic gene expression and cellulose deconstruction [66]. We observed that under normal temperature (25°C) the *hsp30*_0.5kb_:*clr-2* strain failed to grow in liquid cultures containing Avicel, whereas it showed normal growth when sucrose was present instead (Figure S5, S6). Such lack of growth, or the negligible levels of secreted proteins of this strain in Avicel, was comparable to a *Δclr-2* strain (Figure 5, Figure S7). Nevertheless, when Avicel cultures of *hsp30*_0.5kb_:*clr-2* strains were exposed to daily heat-shock pulses, both growth and secreted protein levels were recovered (Figure 5, Figure S7). On the other hand, growth on sucrose was not compromised for either type of strain (Figure S5, S6). Thus, these results highlight the tight regulation provided by this type of promoter, and since both the intensity and frequency of the heat shock treatments can be modified providing a broad and graded response (see Figure 4), it opens up the possibility to maximize, at will, cellulase production in *Neurospora*, likely minimizing fitness costs of CLR-2 overexpression [65]. Notably, although the repeated application of heat shocks has a perceptible effect on mycelial growth, as measured in the WT strain (Figure S8)., the strong induction of a gene of intertest (or in this case the production of cellulases) may fully compensate for growth differences.

**Figure 5.**
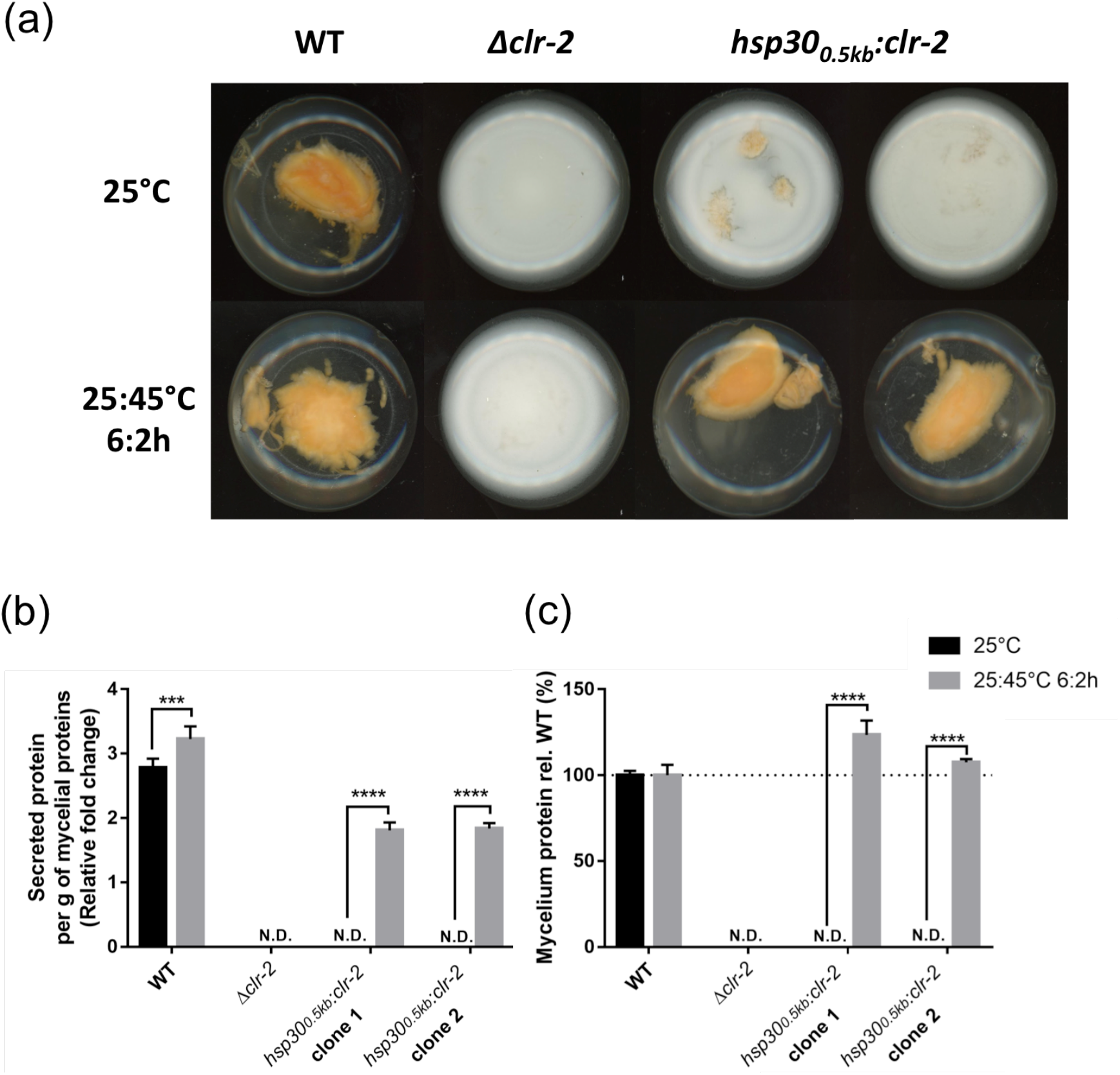
The *hsp30*_0.5kb_ promoter can control a metabolic pathway of biotechnological interest. (**a**) Conidia (10^6^) from WT (x654-1), *Δclr-2, hsp30*_*0*.*5kb*_*:clr-2* (biological clones 1 and 2) were inoculated in Vogel’s media with crystalline cellulose (Avicel, 2%w/v) as carbon source. The flasks were grown in constant light conditions (LL) at 25°C with or without a high-temperature treatment (a pulse at 45°C for 2 h every 6 h; 25:45°C 6:2 h). Before imaging, all the flasks were placed for 7 days in a shaker (125 rpm). The image depicts a representative phenotype of three independent experiments. (**b**) Supernatant protein concentration and (**c**) total mycelial protein content were determined from 7 days cultures of WT, *Δclr-2*, and *hsp30*_*0*.*5kb*_*:clr-2* strains grown on 2% Avicel with or without the heat-temperature treatment (25:45°C 6:2h) as explained in Material and Methods. The supernatant concentration was normalized to the total of mycelial proteins per condition. The mean and standard deviation represent three independent measurements, and three independent experiments. N.D. Not Detected. Statistical significance was performed using a two-way ANOVA plus Sidak’s test (*** = p < 0.001; **** = p < 0.0001).

### Design of synthetic promoters derived from hsp30 putative cis-elements

To generate a versatile molecular tool to be used in *N. crassa* where we could combine the strength and tunable aspects of *hsp30* promoters in a modular fashion, while also reducing the incidence rate of RIP-based mutations, we proceeded to design *hsp30*-based synthetic promoters (*SP30*) using the putative *hse* sequences present in the *hsp30* regulatory region. For this, we selected the predicted minimal promoter region of *hsp30* (up to - 250 bp) and added to it the putative *hse* sequences. In addition to the four predicted *hse* in the *hsp30* promoter (Figure 1A), we identified through bioinformatics tools (see Material and Methods) a fifth *hse* element in *hsp30* (Table S3, S4) with similar p-values to the putative *hse1* and *hse2* sequences, which are the most conserved in *hsp30*. Thus, we designed two short versions of these *hsp30* synthetic promoters: one with 24 bp (*SP30A*) (Figure 6a, Figure S9), and another one with 50 bp spacers between the five *hse* (*SP30B*) (Figure 6b, Figure S9). The synthetic promoters were evaluated *in vivo* by luciferase reporters analyzing their expression profiles over a temperature gradient during long (60 min) and short (1 min) heat pulses (Figure S10). As we envisioned, the two synthetic promoters displayed strong responses to heat shock treatments, showing tunability to variation in both temperature and exposure times (Figure 6c,d).

**Figure 6.**
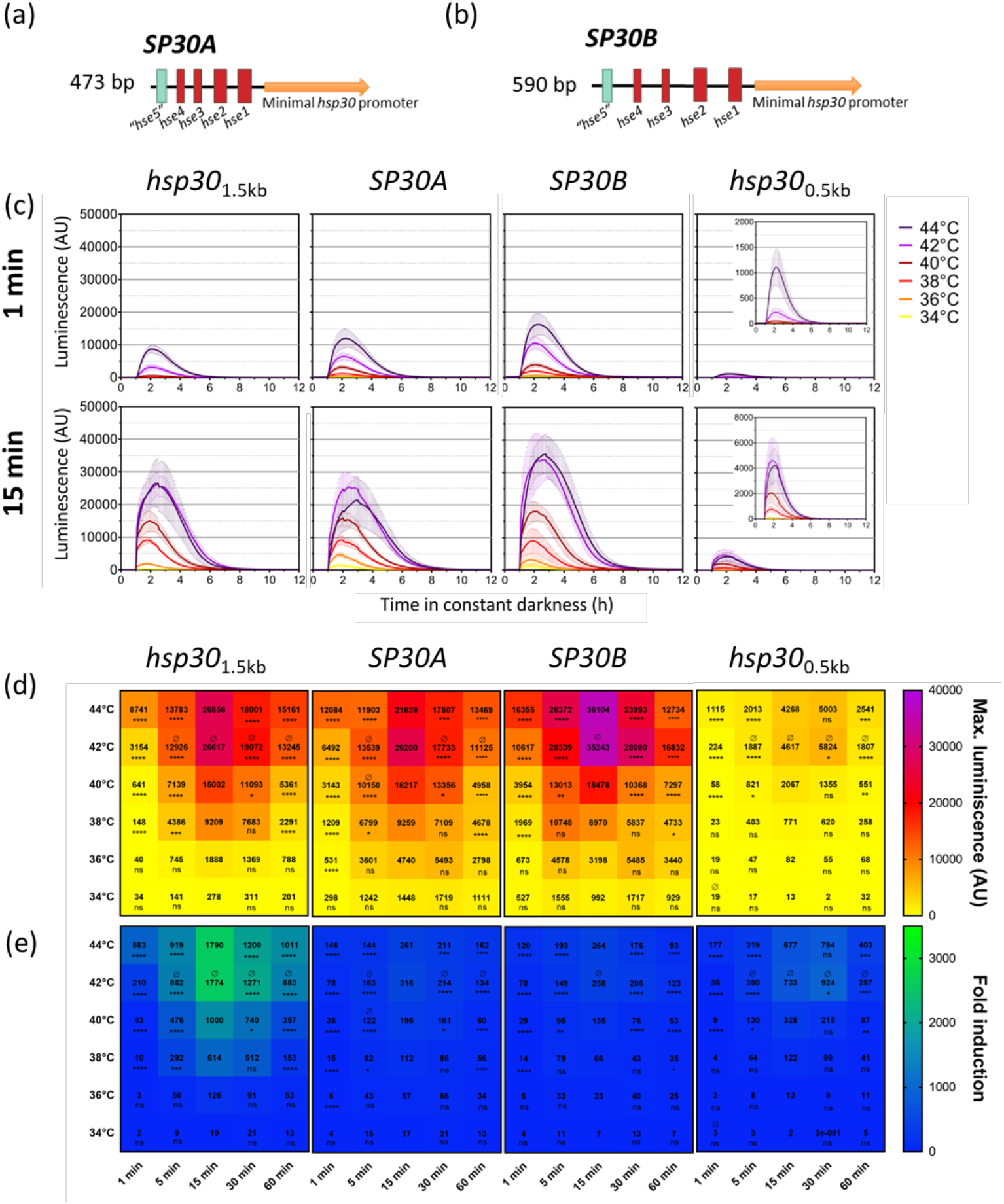
Design of two inducible synthetic promoters tunable by subtle changes in heat shock treatments. (**a** and **b**) *hsp30* synthetic promoters (*SP30*) scheme, where synthetic promoters with (**a**) 25 (*SP30A*) or (**b**) 50 bp (*SP30B*) spacers between the indicated putative *hse* were used to generate reporter genes in the context of a minimal *hsp30* promoter of 250 bp. (**c**) Luciferase activity profiles of the synthetic promoters against a temperature gradient for 1 min and 15 min. A close-up of the *hsp30*_0.5kb_ graph is displayed on the left side. Each curve corresponds to the average of two or three biological clones with four independent wells each ± standard deviation (SD), and represents the behavior in two independent experiments. (**b**) Maximum luminescence and (**c**) fold-changes observed after the heat-shock treatments for the indicated *hsp30* promoters are indicated. The maximum luminescence was defined as the average of the highest values. Fold induction was calculated based on the maximum luciferase expression (shown in panel **b**) with respect to the average of the background values before heat shock treatment of each promoter region. The data was obtained from the luciferase activity profile shown in Figure S10. Statistical significance was performed using a two-way ANOVA plus Dunnett’s test (for time treatments all values were compared to 15 minutes: *= p<0.05; **= p < 0.01; *** = p < 0.001; **** = p < 0.0001; ns = non-significant. For temperature treatments all values were compared to 44°C, where Ø indicates non-significant. Time significance is indicated below each value, whereas for temperature only non-significance (Ø) is shown above the value when it corresponds.

While the native *hsp30* promoter has an outstanding ON/OFF ratio, given in part by the extremely low basal levels of expression, the synthetic promoters displayed higher basal activity, which negatively affects such relationship (Figure 6e, Figure S11). Nevertheless, the basal levels of these promoters are considerably lower than the ones of *hsp70* and *hsp80* (Figure 2, Figure S1). Both synthetic promoters behave similarly, exhibiting (across all the temperatures/exposure times), equivalent or higher maximal activity than the native *hsp30* system (Figure S10). Despite this high activity upon induction, due to their increased basal expression, they yield less fold of induction compared to *hsp30*_0.5kb_ (Figure S12).

Thus, these synthetic *hsp30* promoters represent a viable strategy to provide heat-shock response of genes of interest, in a modular fashion, and conserving the key characteristics of the native *hsp30* system regarding. tunability and maximum activity.

## Discussion

Changes in temperature are a ubiquitous physical stimulus to which all organisms are exposed to and, consequently, they have developed mechanisms that help them cope with strong increases in temperature such as heat shocks. These mechanisms involve heat shock proteins (HSPs) and their accurate transcriptional regulation, which play a relevant role in high-temperature tolerance. In addition to its protective function, it has been possible to use *hsp* promoters as molecular tools for spatial-temporal control of genes of interest in several organisms [39, 67–69], strategy that has been poorly exploited in filamentous fungi. In this work, we profiled the response of three *N. crassa hsp* promoters (and their resected versions) to discrete changes in temperature for different treatment duration, to obtain a better understanding of the dynamic biological response of such genes. Moreover, we aimed to select the most suitable candidate to be used as an inducible system that, more than just an ON-OFF switch, could act as a rheostat to provide graded transcriptional responses, depending on the intensity of the physical stimuli.

All the herein characterized Neurospora promoters (*hsp30, hsp70, hsp80*) showed a rapid and strong response under standard heat shock treatment in an incubator: 45°C for 1 hour [70], being the *hsp30*_1.5kb_ the one with the highest one. It is known that low molecular weight HSPs tend to be the first ones to act against protein denaturation at high temperatures, due to their inability to bind and hydrolyze ATP, like other larger HSPs, and would have a strong response to heat stress [71]. Therefore, the higher response levels of the *hsp30* promoter appears to correlate with their described role upon heat stress. Expression levels observed under non-inducing conditions in this work revealed a basal constitutive expression of the *hsp70* and *hsp80* genes in contrast to *hsp30*, which goes from extremely low expression levels to strong induction in response to thermal stimulation. Previous investigations reported that *hsp30* mRNA levels are rather negligible at normal growth temperatures [72], while high basal expression is characteristic of the HSP members of the HSP70 and HSP90 families [73].

The resected version of these promoters provided information on the possible roles of their relevant *cis*-elements. In *hsp70*, these elements are located in the *hsp70*_0.6kb_ region, and the absence of further upstream sequence did not cause a decrease in the thermal response. On the contrary, the relevant regulatory elements in *hsp80* appear to be upstream of the *hsp80*_0.6kb_ region, because this resected section failed to show induction despite having the majority of the proposed *tre* like boxes. In contrast, the *hsp30*_1kb_ region had the same tunability and responsiveness as *hsp30*_1.5kb_, suggesting that several of the relevant elements are already present in this resected region. On the other hand, the presence of only two out of the four proposed *hse* in *hsp30*_0.5kb_ may explain its strong activity at 45°C, but diminished responses closer to 35°C, as it has been reported that having multiple *hse* tends to confer higher response levels [73,74].

Our data indicate that the analyzed *hsp30* sections display key features of an ideal regulable promoter, allowing progressive control of transcription at different stimuli intensities. It also provides tight regulation that can maintain low basal levels of expression in the absence of thermal stimulation, in addition to a rheostat-like behavior (high tunability) and temporal control [75]. The *qa-2* and *vvd* promoters are probably some of the most used inducible promoters in *Neurospora*; however, *qa-2* has a limited range of expression, and a low, albeit rather leaky basal transcription in the absence of Quinic Acid [21]. It also has the additional caveat that once the inducer is added, it is cumbersome to remove it, as it would require media exchange. Overall, the *vvd* and *hsp30* promoters share key properties as, for example, no need to supplement the growth media with an inducer as they both respond to a physical stimulus that can be externally and easily provided [75]. However, the particular light/dark condition requirements for controlling *vvd* [18] imply defined lab settings (dark-room), and its use may not be fully compatible with photobiology or circadian studies.

Thus, the described *hsp30* system shows flexibility as the response can be fine-tuned not only by the inducing temperature but also by the length of the treatment (Figure 4; Figure S4). This was clearly exemplified in the discrete changes in temperature, provided in the thermocycler approach, where the selected reporter strains were treated at six different high temperatures simultaneously. Such simple experimental setup for heat shock treatment provides several advantages such as: *i)* It allows the exposure of the strains to a wide range of temperatures; *ii)* Optimizes the time required between experiments; *iii)* Increases the precision and accuracy of the temperature at which the strains are exposed (particularly as PCR tubes are designed for efficient heat transmission). Although with this we were able to obtain an accurate profile of the *Neurospora hsp30* promoter responses, the limitations of the used methodology should also be noted. The activity profiles at high temperatures for prolonged treatments (60 min and 30 min) can be weaker than the ones obtained at shorter times, which is probably caused by the efficient temperature transfer in the thermocycler compared to the one inside an incubator, where the former might result - at longer time points-in high stress compromising cellular status, impairing reporter output. In addition, the use of 96-well format (PCR-tubes) limits the amounts of utilized media, restricting growth as well. In our hands, the tested strains grew well for at least 2 full days.

The map of *hsp30* responses to different temperatures shows a particular pattern: at short exposure times, luciferase expression mostly correlates with the increasing temperatures, and also there is a clear gradient of responses between 15 min and 1 min treatments (Figure 4b; Figure S4). Nevertheless, at longer exposure times, a diminished response is generally observed if compared to 15 min exposures. As mentioned earlier, this could be attributed to excessive stress, caused by the direct and prolonged exposure to high temperatures, which could also explain the augmented dispersion of the data obtained at such temperatures and exposure times. Despite this, we could observe a wide range of response intensities depending on the degree of the heat shock and the duration of this stress for the *hsp30* promoters (Figure 4). Importantly, the data obtained herein reveal that the latter are highly modulable in a range of high temperatures, exhibiting also an analogue (gradual and continuous) and not digital/binary (ON-OFF) [76] response.

As proof of principle, we conditioned *Neurospora* cellulolytic capabilities, by putting *clr-2* expression under the control of the *hsp30*_0.5kb_ promoter. Notably, although the resected promoter that we used has lower activity levels compared to longer *hsp30* sections, the results clearly demonstrate how our system is capable of tightly regulating a gene of interest that can have ecophysiological implications [77] and industrial impact [78]. A main obstacle in the industrial application of the degradation of cellulose has been the high cost of enzyme production, which has restricted accessing lower prices of bioethanol as a fuel alternative, a discussion that is revived every time gas prices are on the rise [79,80]. Using a *hsp30*_0.5kb_ promoter and temperature treatments to regulate *clr-2* expression, we reverted the poor growth phenotype in cellulose (equivalent to the one seen in a *Δclr-2* strain) although secreted proteins levels were lower compared to a WT strain (Figure 5). Nevertheless, when we used a different protocol, consisting in transferring sucrose-grown mycelia to Avicel, we observed low levels of protein secretion for the *hsp30*_*0*.*5kb*_*:clr-2* strains at 25°C, situation that was reverted - surpassing even WT levels - when heat pulses were applied (Figure S7). In addition, other induction protocols (with more frequent heat shocks), or implementing other *hsp30* promoter synthetic versions could easily yield augmented cellulase levels.

While further analyses are necessary to advance and optimize this methodology, the thermal induction strategy herein described presents itself as an attractive alternative to regulate the expression of genes of interest and to tightly regulate and tune desired phenotypes.

Furthermore, to facilitate the adoption of a *hsp30*-based system, we designed a synthetic version of *hsp30*, for which we utilized the *hsp30*_1.5kb_ promoter’s putative *hse*. Despite the lack of nucleotide-resolution studies characterizing these *hse* as functional, their sequence identity strongly suggests that these regions are conserved and are likely recognized by HSFs, commanded by the major regulator in *Neurospora*: HSF-1 (NCU08512). In addition, it has been shown that the HSF of *Neurospora* can efficiently recognize a *hse* from yeast [81]. Based on this, we used one of the HSF-1 motifs obtained from yeast and *N. crassa* to further identify new, and confirm the previously proposed *hse* in the *hsp30* promoter (Tabla S2, S3). Thus, we detected a new *hse* (*hse5*) that may also play a part of the *hsp30*_1kb_ regulation. With this information, we designed two synthetic promoters based on the five putative *hse* present in *hsp30*, where we contemplated having different lengths of spacers between the *hse*. It has been observed that the optimal spacer between the *cis*-elements can maximize response, although such levels would depend on promoter/organismal context. In bacteria the optimal distance between the minimal *cis*-elements in the core promoter is 17 bp for *E. coli* [82], but with higher lengths up to 80 bp as in the case of Pseudomonas [83]; whereas in eukaryotes the minimal distance between TATA box and the transcriptional start site (tss) are ∼30bp, observed in yeast [84] and mammals [85].

We were able to generate a high range of tunability, although the low background expression of the native *hsp30* promoter was not fully maintained (Figure 6, Figure S1, S11). Indeed, issues like this can be sometimes a tradeoff of synthetic promoters, where despite strong responses basal expression is higher than expected [86]. Nevertheless, reproducibility, tunability, and temporal controllability are properties that are still present in the designed synthetic *hsp30* promoters. In addition, the high conservation in the regulation of *hsp* expression allows predicting that some of these resected promoters could readily work in other ascomycetes (Figure S13). In particular, it will be interesting to attest the behavior of the modular synthetic *hsp30* promoters in biotechnologically relevant fungi such as *Aspergillus niger* or *Trichoderma reesei*.

Thus, in this work we provided a detailed profile of the response of the *hsp* genes to thermal stimuli, while also extending the molecular tools available for *N. crassa* by describing new set of inducible heat shock promoters with overall low background levels, and allowing a rheostat-like adjustment of expression of a gene of interest.

## Materials and Methods

### Plasmid Construction

All the plasmids were assembled by yeast recombinational cloning [87] in *Saccharomyces cerevisiae*, strain BY4741 (*MATa, his3Δ1, leu2Δ0, met15Δ0, ura3Δ0*), amplifying the promoter fragments from WT (74A) *N. crassa* genomic DNA. For the synthetic promoters *SP30A* and *SP30B*, the fragments were synthesized by GENEWIZE (https://www.genewiz.com/) and then cloned as described above. The information of backbone plasmids, PCR products and primers used for each construction are detailed in Table S1. All the constructions generated were confirmed by sequencing.

### Strains and culture conditions

The transcriptional reporter strains, containing the analyzed promoters (full, resected, and synthetic) were fused to a destabilized luciferase (*luc*^_*PEST*_^) and targeted to the *csr-1* locus for cyclosporine selection [14], and transformed in a selected strain (x654-1: *ras1*^*bd*^, *mus51*^*RIP*^, *a*) as previously reported [88], following a standard electroporation protocol [11].

The *Δclr-2* strain (xc2386-2; *Δclr-2, ras1*^*bd*^, *mus51*^*RIP*^, *a*) was obtain from the cross between #15834 (A) and L418T654c-1 (*ras1*^*bd*^, *mus51*^*RIP*^, *a*). The *hsp30*_0.5kb_:*clr-2* (xc2417; *ras-1*^*bd*^, *mus51*^*RIP*^) strains used for cellulose phenotypic assays were obtained by replacing the 2000 bp upstream region (relative to the ORF) of the *clr-2* gene (NCU08042), with the *hsp30*_0.5kb_ promoter in a x654-1 background (primers on Table S1). The selection was made by incorporating a *bar* resistance cassette upstream the resected promoter, and homokaryotic strains were obtained through sexual crosses [89].

The vegetative growth utilized slants with 1X Vogel minimal medium (VM) [90] supplemented with 2%w/v sucrose in 1.5%w/v agar, for 5-7 days in constant light (LL) at 25°C, whereas for sexual crosses synthetic crossing medium (SCM) [89] was used. Sorbose-containing medium (FIGS) was utilized for colony isolation and ascospore germination [91]. Ascospores were picked on slants containing VM media supplemented with bialaphos [92], cyclosporine (5 µg/mL), and/or luciferin (10 µM), in order to select for progeny carrying knockout cassettes and/or reporter activity. To conduct luciferase analyses in both Heat Shock treatment setups (see below, 2.3) we used LNN-CCD media (0.03% glucose, 0.05% arginine, 50 ng/ml biotin, 1.5% agar) [93] supplemented with the indicated concentrations of luciferin.

### Heat Shock Analysis

The Heat shock treatments were conducted applying two different strategies: first using an incubator for exploratory functional analyses; and second, using a gradient thermocycler for high-throughput and more accurate studies.

#### a) Incubator

The strains were grown in black 96-deep well cell culture plates for 24h in LL at 25°C with 750 µL of LNN-CCD media and luciferin (0.5 mM) per well, and the plates were covered with a breathable transparent membrane. After 24h of monitoring LUC activity, a temperature pulse was provided during one hour of treatment. The temperatures chosen for this exploratory analysis were: 30°C, 35°C, and 45°C utilizing a Percival incubator.

#### b) Thermocycler

The strains were inoculated in 96-well plates made of PCR tubes, covered outside with black spray paint resistant to temperature (Rust-Oleum) to avoid light cross-contamination, and covered with a breathable transparent membrane. Strains were grown for 5 h in LL plus 12h at constant darkness conditions (DD) at 25°C, using 50 µL of LNN-CCD media and luciferin (0.5 mM). After that, the LUC activity was monitored for one hour to then apply a temperature treatment exposing the strains to a temperature gradient (34°C, 36°C, 38°C, 40°C, 42°C, and 44°C) of variable treatment times (60 min, 30 min, 15 min, 5 min, or 1 min) in a thermocycler (Veriti™, Applied Biosystems™ 4375786) to finally record the LUC activity for the following 12h. A scheme of this heat shock treatment methodology can be found in Figure S1. Importantly, the hot lid of the equipment was manually disengaged in order to eliminate additional sources of heat that could interfere with the induction protocol. Both strategies then imply the use of Percival incubators equipped with CCD PIXIS 1024B cameras (Princeton Instruments) to register the luciferase expression using acquisition settings of 5 minutes of exposition and 3 or 12 frames per hour to incubator and thermocycler strategy, respectively [93].

### Comparison of Neurospora crassa inducible promoters

We compared, using luciferase transcriptional reporters, *hsp30* with the well-known light-inducible promoter *vvd* (NCU03967; 3.5 Kb upstream region) and the Quinic acid (QA) inducible promoter of *qa-2* (NCU06023; 600 bp upstream region). The primers and plasmids used to generate these constructions are detailed in Table S1. The analyses were conducted in 96-deep well cell plates and PCR tubes using LNN-CCD media with luciferin as detailed above. To compare light, temperature, and QA induction, strains were grown overnight at 25°C in LL, and then LUC activity was monitored at 25°C in DD. After 5 hours, strains were subjected to the corresponding treatment:

a. Light pulse: One hour at 25°C of white light (100 µM/m2/s; wavelength 400–720 nm).
b. Temperature pulse: One hour at 45°C in DD, depending on the experimental set up the heat pulse was given using a thermocycler or an incubator as indicated previously.
c. QA induction: A drop of QA 1 M was added to each 96-well tube, to obtain a final concentration of 0.01M, after which LUC activity was immediately monitored.

### Growth on cellulose phenotypic assays

Flasks containing 50 mL of minimal Vogel’s 1X media supplemented with 2% w/v sucrose or Avicel were inoculated with conidial suspensions. Strains were grown for seven days in agitation in Percival incubators in LL, where control strains were kept at 25°C and strains subjected to temperature treatments received a temperature pulse of two hours at 45°C every eight hours (three times a day) for 7 days.

Additionally, flasks containing 50 mL of minimal Vogel’s 1X media supplemented with 2% w/v Sucrose were inoculated with conidial suspensions. Strains were grown for 48h in agitation in Percival incubators in LL, then mycelia were washed and transferred to flasks containing 50 mL of minimal Vogel’s 1X media supplemented with 2% w/v Avicel, where control strains were kept at 25°C and strains subjected to temperature treatments received a temperature pulse of two hours at 45°C every eight hours (three times a day) for 24 h. Biomass and protein quantification of each condition was performed using dry-weight of grown mycelium and Bradford curve interpolation, respectively as previously described [9].

### Heat shock element (hse) sequence analysis

We used the YeTFaSCo database (http://yetfasco.ccbr.utoronto.ca/index.php) for *S. cerevisiae* HSF-1 based motifs. The motif ID 615 was used as a matrix to identify new *hse* or to confirm the previously described *hse* in the *hsp30* promoter using FIMO of MEME suite (Version 5.4.1). A p-value < 0.001 was used to select the *hse*. The results of this analysis can be found in Table S2.

For the *hse* identification in *hsp30* promoter through the HSF-1 motif of *N. crassa*, we used the CIS-BP database (http://cisbp.ccbr.utoronto.ca/) [94]. The motif ID T243453 (HSF-1, NCU08512) was utilized to analyze the *hsp30* promoter sequence using the tool “Scan single sequences for TF binding” (Motif model: PWMs-LogOdds). The default threshold was utilized (score not under 8). The results of this analysis can be found in Table S3.

For the *hse* alignments, the promoters of *hsp30* in fungal orthologs (*N. crassa* NCU09364, *N. tetrasperma* NEUTE1DRAFT_72918, *N. discrete* NEUDI_159228, *T. reesie* TRIREDRAFT_122363, *A. niger* M747DRAFT_254277, *A. nidulans* AN2530, *A. fumigatus* Afu3g14540) were all downloaded from FungiDB and the DNA sequence alignment was performed with MEGA version 11 software with ClustalW algorithm [95]. The putative *hse* sequences were sought individually at each promoter and then, with these sequences, multiple nucleotide-sequence alignments were performed as indicated above.

### Statistical analysis

Graphs and statistical analyses were made using GraphPad (Prism) version 7.0.

## Acknowledgements

This work was funded by ANID—Millennium Science Initiative Program—Millennium Institute for Integrative Biology (iBio ICN17_022), ANID/FONDECYT 1211715, the International Research Scholar program of the Howard Hughes Medical Institute, and The Richard Lounsbery Foundation. Additonal funding was provided by ANID/FONDECYT Postdoctorado 3220747 to A.G. and ANID/FONDECYT Postdoctorado 3220597 to F.M.G.

