## Supplemental Figures and Tables for "Developing a temperature-inducible transcriptional rheostat in *Neurospora crassa*"

**Supplemental Material**

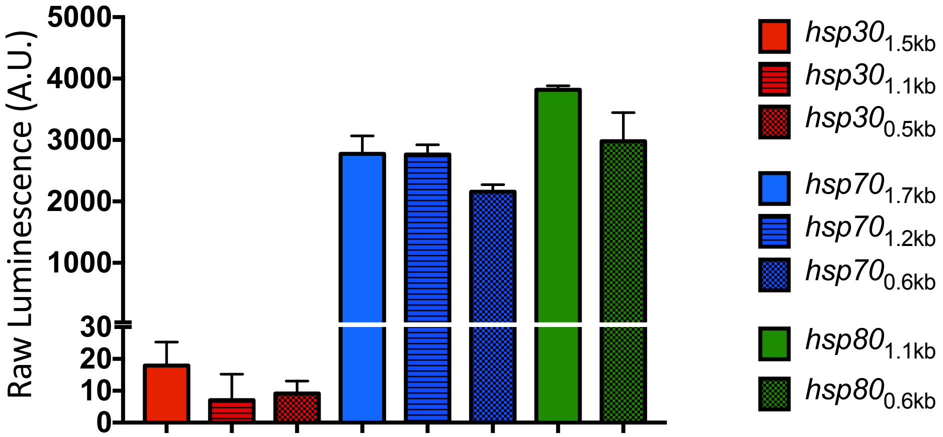


**Figure S1. Basal activity of *hsp* reporters.** Basal (background) luminescence levels of each *hsp* promoter and their resected sections. Each bar indicates the average of two or three biological clones with four independent wells each ± standard deviation (SD), and represents the behavior of two independent experiments. Values were obtained prior to delivering the heat shocks, in a 96-well plate format.

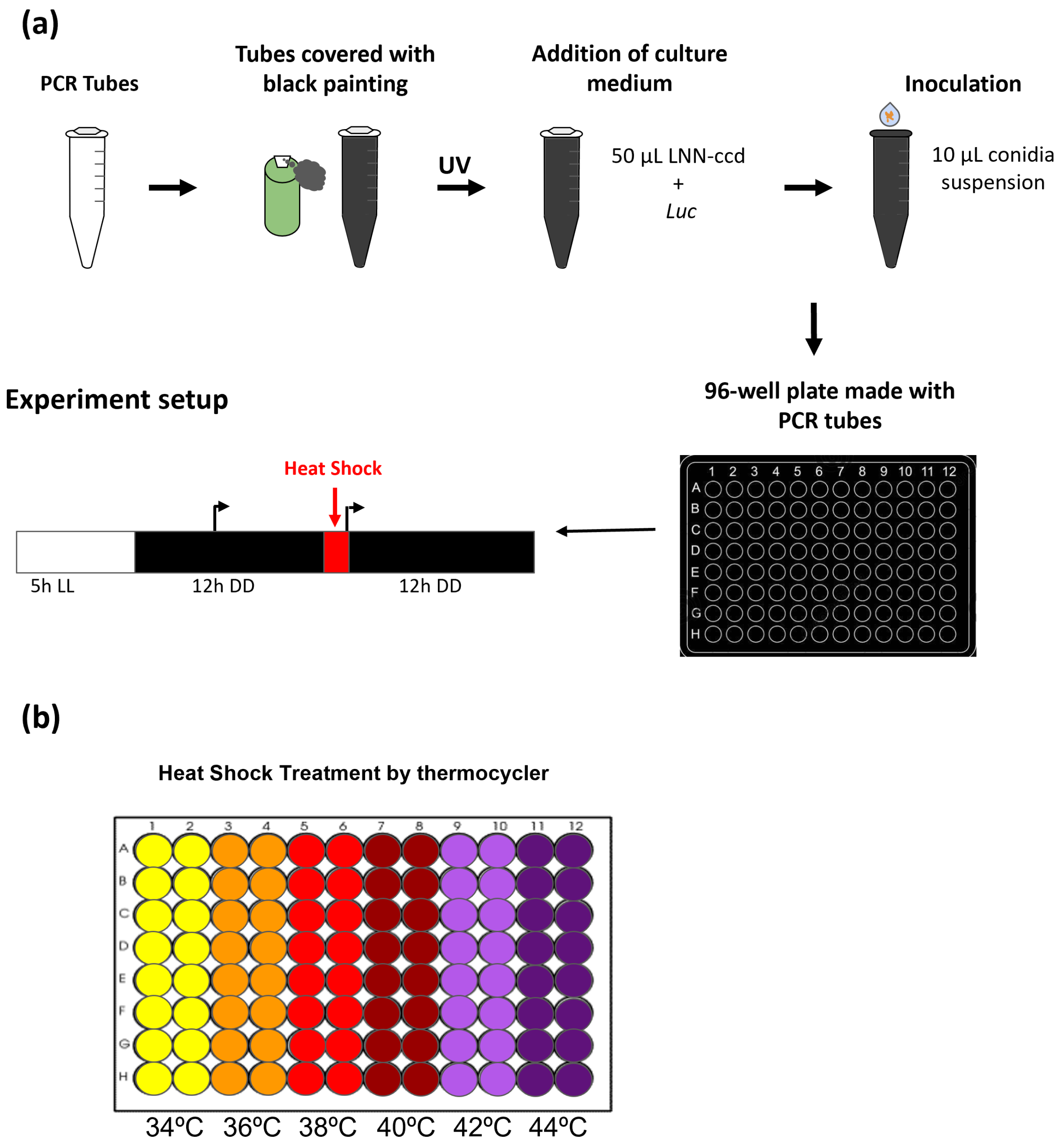


**Figure S2. Strategy to expose strains to heat shock in a temperature gradient.** (**a**) Scheme of the methodology utilized to generate a darkened 96-well plate with PCR tubes. The PCR tubes were externally painted with black aerosol, and then sterilized with UV light for 15 min. LNNccd media with luciferin (0.5 mM) was added, and then the strains were inoculated as conidia suspensions. The 96-well plate was placed in constant light (LL) at 25ºC for 5 h, and then transferred to DD for 12h at 25ºC. Background luciferase levels were calculated for 1 hour prior to the heat-shock. Luminescence was acquired with a CCD camera (indicated with broken arrows in the "Experimental setup" diagram), and tubes were exposed to heat treatment (in a gradient thermocycler) for different times (60, 30, 15, 5, and 1 min). We continued measuring luminescence levels after the heat shock for 12 additional hours. (**b**) Scheme of the temperatures used for the heat- shock treatment in the gradient thermocycler in a range of 34ºC to 44ºC.

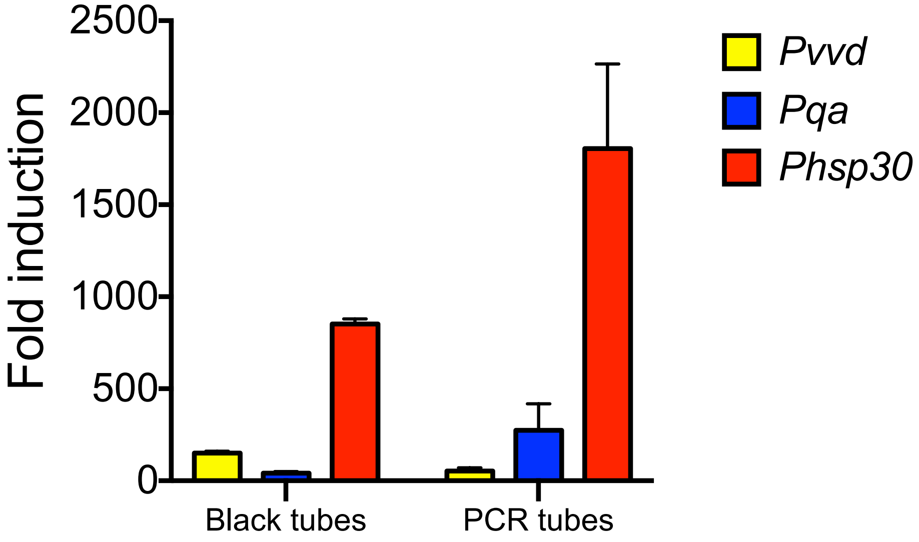


**Figure S3. Comparison of fold-changes achieved by *hsp30* and other inducible *N. crassa* promoters**. Fold induction achieved by the *hsp30, vvd* and *qa (qa-2)* promoters after providing the respective cognate stimuli. Fold induction was calculated with the maximum luciferase expression (the average of the three highest consecutive values) respect to the average of the background values before inducing each promoter.

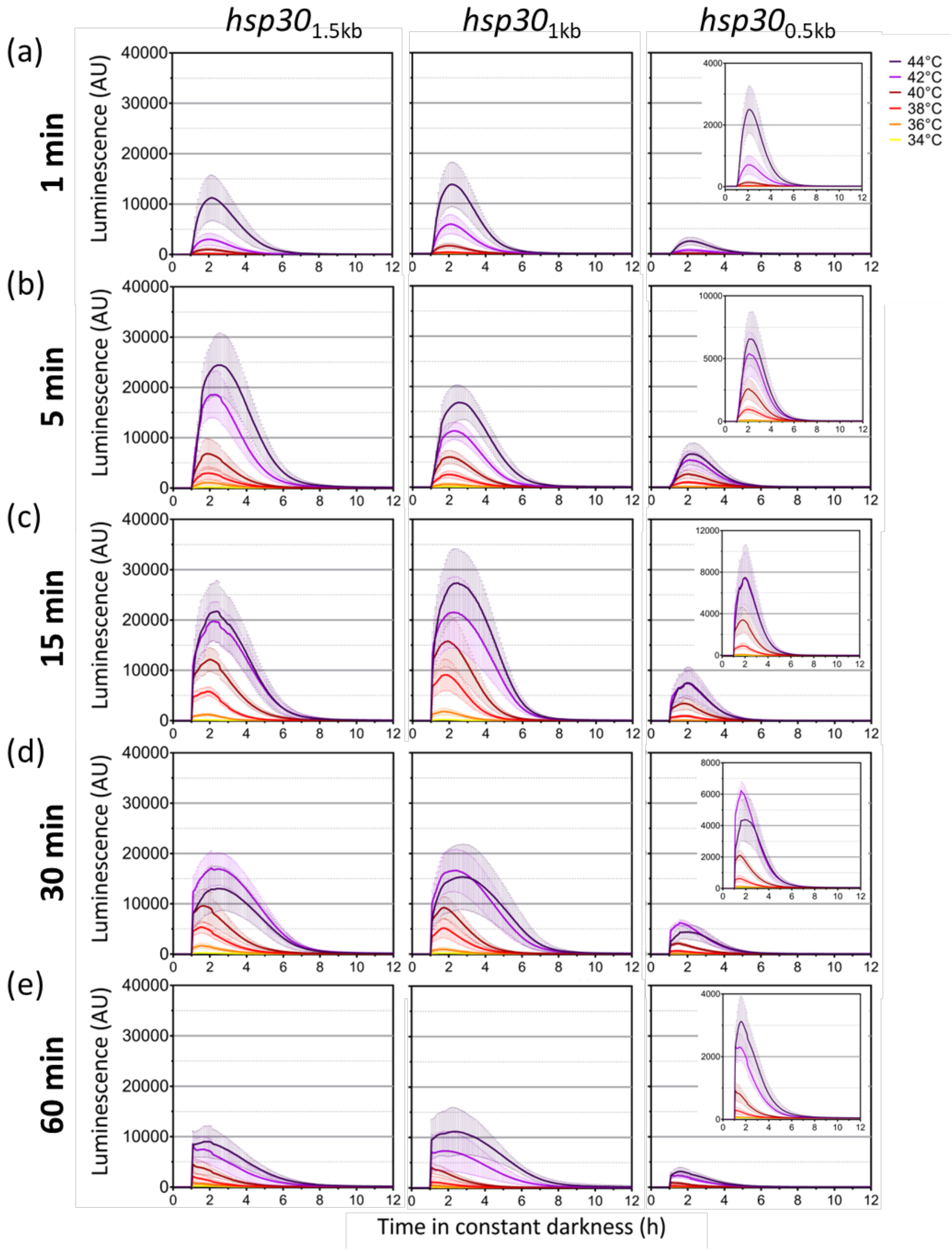


**Figure S4. Luciferase activity profiles conferred by full or resected *hsp30* promoters to a temperature gradient and different exposure times**. (**a** to **e**) Activity profiles of each *hsp30* promoter region after a short (**a** to **b**) or long (**c** and **e**) heat shock treatment. Average and SD of each measurement are shown (2 biological clones, with eight technical replicas for each one). A close-up of the *hsp30*$_{0.5kb}$ graph is displayed up on the right side when needed. The methodology is described in Figure S2.

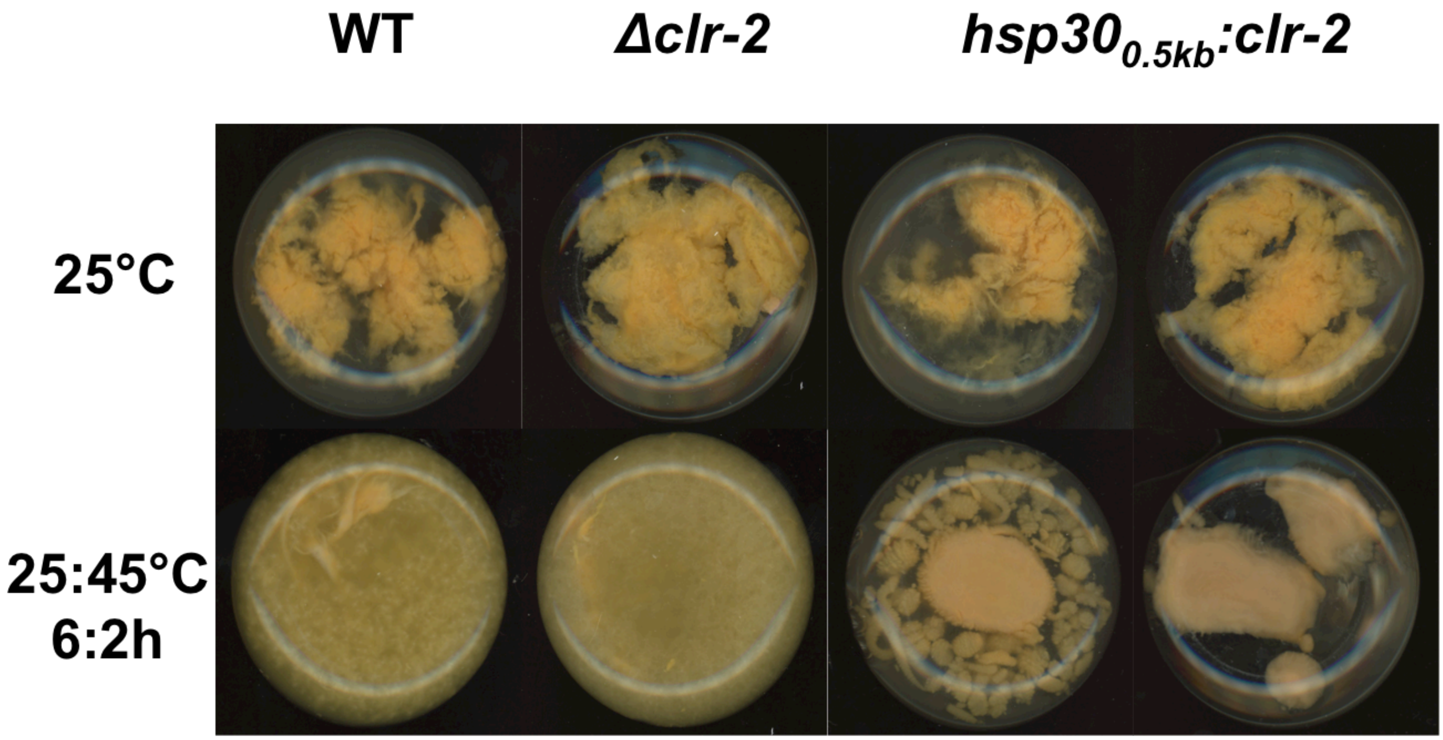


**Figure S5. Phenotypic analyses of heat shock treatments in sucrose media.** Conidia ($10^6$) from WT (x654-1), *Δclr-2*, *hsp30*$_{0.5kb}$*:clr-2* (biological clones 1 and 2) were inoculated in Vogel's media with sucrose (2%w/v) as carbon source. Flasks were grown in constant lights (LL) at 25°C with or without a high-temperature treatment, the latter corresponding of a 45°C pulse for 2 h every 6 h (25:45°C 6:2 h). Cultures were kept for 7 days in a shaker (125 rpm). The photographs are representative of three independent experiments.

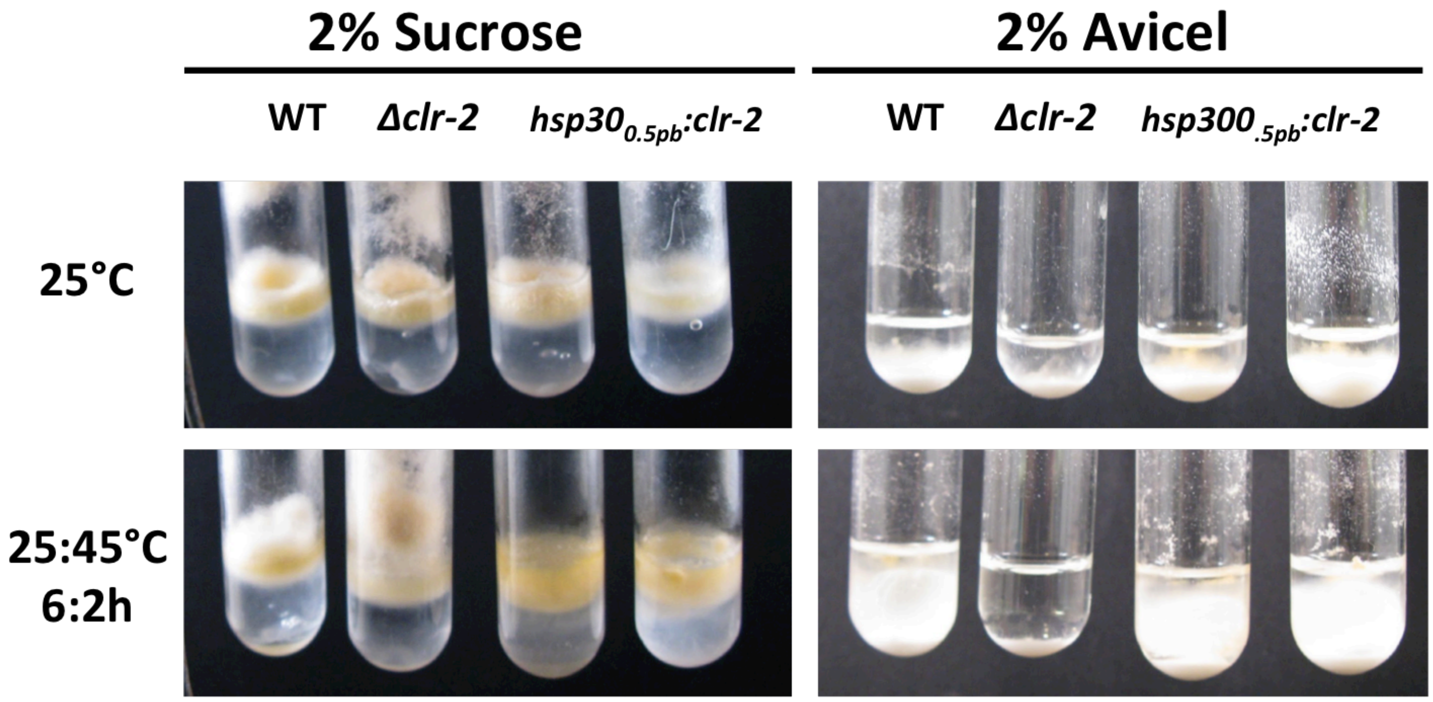


**Figure S6. The *hsp30*0.5kb promoter can control a catabolic pathway of biotechnological interest.** Conidia ($10^6$) from WT (x654-1), Δ*clr-2*, *hsp30*0.5kb:*clr-2* (biological clones 1 and 2) were grown in Vogel's media with sucrose (2%w/v) and crystalline cellulose (Avicel, 2%w/v) as carbon source. One set of tubes grew at 25ºC, while the others were exposed to a cycle of 45ºC for 2 h and 25ºC for 6 h (repeated 3 times every 24 h). All tubes were placed in a shaker (125 rpm) in constant lights (LL), for 4 days (sucrose) or for 7 days (Avicel). The photographs are representative of the behavior of three independent experiments.

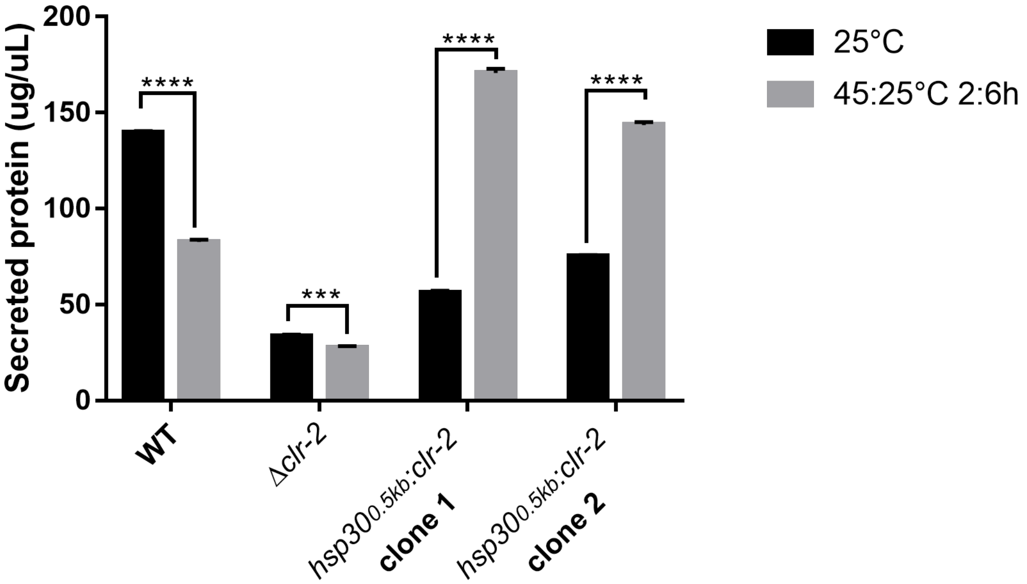


**Figure S7. Secreted protein levels in shifted cultures.** (**a**) Conidia ($10^6$) from WT (x654-1), *Δclr-2*, *hsp30*$_{0.5kb}$*:clr-2* (biological clones 1 and 2) were inoculated in Vogel's media with sucrose and were grown in constant light conditions (LL) at 25°C for 48h. Then mycelia were washed and transferred to Vogel's media with crystalline cellulose (Avicel, 2%w/v) as carbon source, and the flaks were grown in constant light conditions (LL) at 25°C with or without a high-temperature treatment (a pulse at 45°C for 2h every 6h; 25:45°C 6:2h). Supernatant protein concentrations were determined from 24 h cultures of WT, *Δclr-2*, and *hsp30*$_{0.5kb}$*:clr-2* strains grown on 2% Avicel with or without the heat-temperature treatment (25:45°C 6:2h) as explained in Material and Methods. The mean and standard deviation represent three independent measurements, in three independent experiments. Statistical significance was performed using a two-way ANOVA plus Sidak's test (*** = $p < 0.001$; **** = $p < 0.0001$).

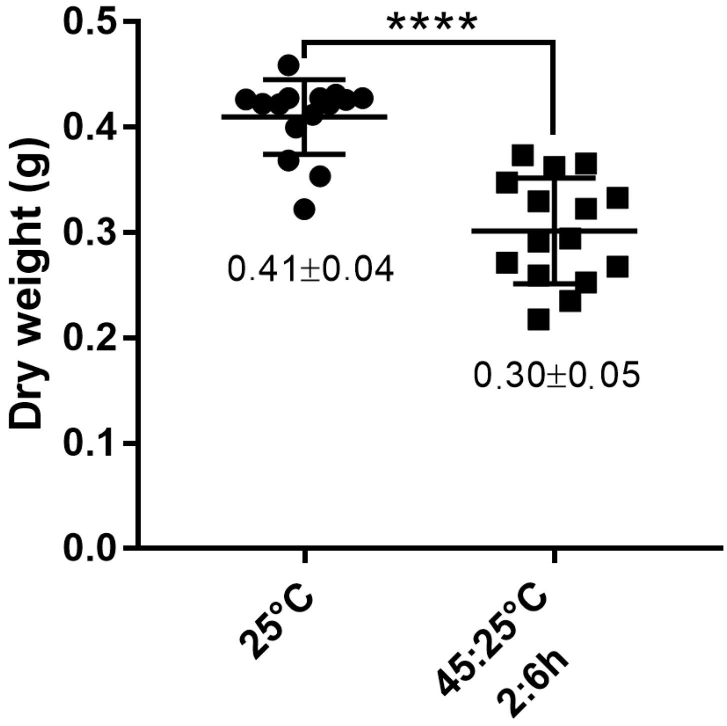


**Figure S8. Effect of high temperature treatment in WT growth.** Conidia ($10^6$) from WT (x654-1) was inoculated in Vogel's media with sucrose (2%w/v) as carbon source. Flasks were grown in constant lights (LL) at 25°C with or without a high-temperature treatment (a pulse at 45°C for 2h every 6h, 25:45°C 6:2h). Cultures were kept for 4 days in a shaker (125 rpm) and then the mycelium was harvested and dried. Statistical significance was determined by a t-student (**** = $p < 0.0001$).

>SP30A

aatacattgatcacctggaaccttcttgaatgctcttcagatcttatatcatctagtaaactagcagtaccaagctatcaaacctttacctcggcgtgaaattaccagaaactgcgaccgggtgcagcgcagtgtggaactttctaggctgctcccgaagcgtgcgattggccagcgaattacagtggccattctagaatgacctggcatctgcatcaaatgctccttccaccttcttcgtccttcgcatcacaggtgaaagacaaggtaatttgtgaagcaaacaatgctcccactcaaatataaatctgggtgtgatgtctcccttttcatattgtcgattctctgttcagcagatcaagatcatccagcaagcgaagtaatcactctgaacactctcaacagcatctactacactcagcaaacgcacagatacctccgtcgccactcttttaacacacctaaagtcaaaa

>SP30B

caaaacgggattcaatacattgatcacctggaaccttcttgaatgctcttcagatcttatacagctactacagtttcataatcagtctcatctagtaaactagcagtaccaagctatcaaaacaaactagttaggaaggaaatattccctttacctcggcgtgaaattaccagaaactgcgaccactgattgatccggcaaaactgaagcgggtgcagcgcagtgtggaactttctaggctgctcccgaagcgtgcgattggcctcacaccgtctccccaagagaactccagcgaattacagtggccattctagaatgacctggcatctgcatcaaatgctccttccaccttcttcgtccttcgcatcacaggtgaaagacaaggtaatttgtgaagcaaacaatgctcccactcaaatataaatctgggtgtgatgtctcccttttcatattgtcgattctctgttcagcagatcaagatcatccagcaagcgaagtaatcactctgaacactctcaacagcatctactacactcagcaaacgcacagatacctccgtcgccactcttttaacacacctaaagtcaaaa

**Figure S9. FASTA sequence of the *SP30A* and *SP30B* promoters.** Sequence of both synthetic promoters (*SP30A* and *SP30B*). *hse* (red), the newly assigned *hse* (light-blue), spacers (black) and the putative minimal promoter of *hsp30* (deep red) are indicated.

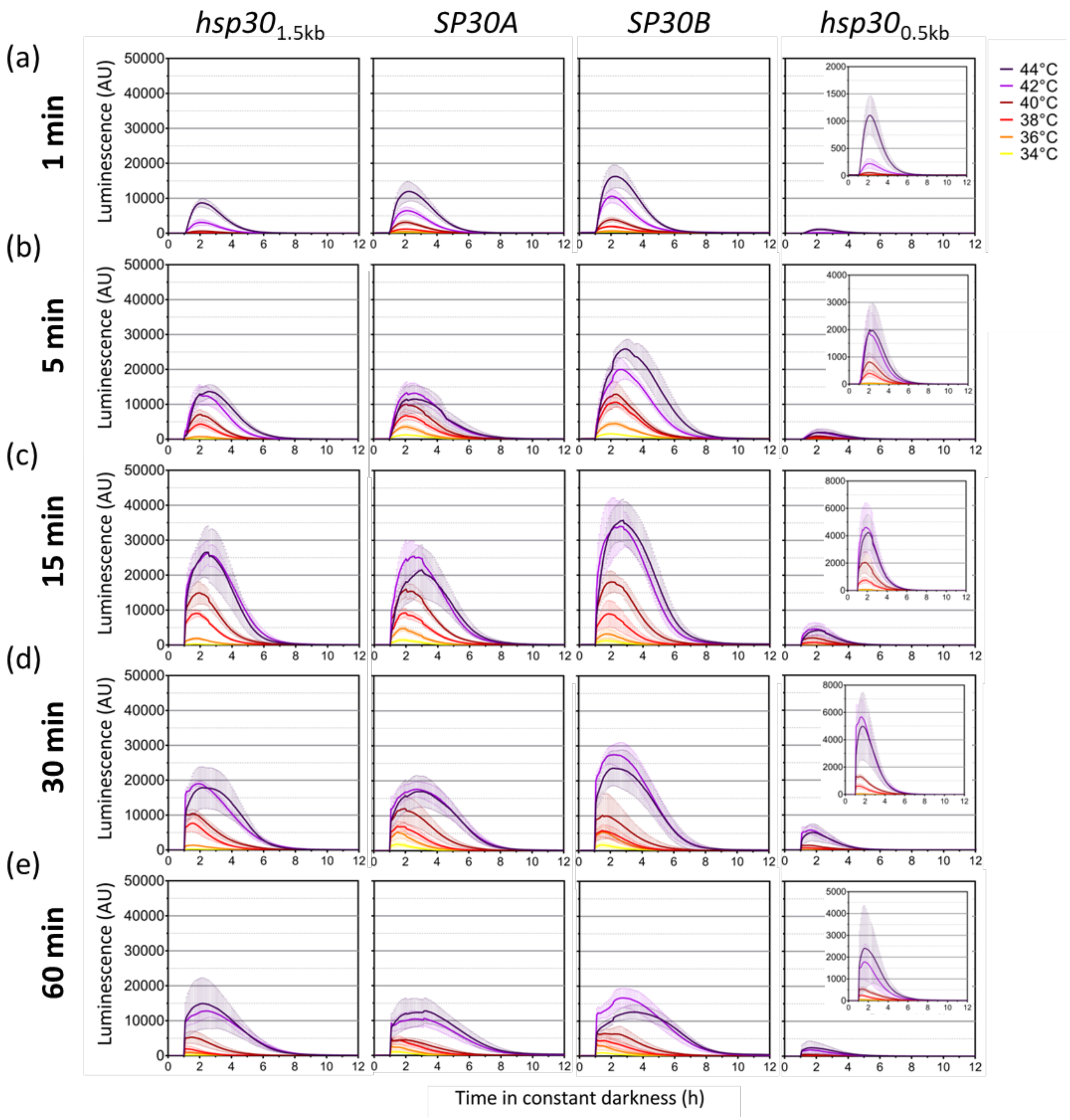


**Figure S10. Luciferase activity profiles conferred by the *SP30A* and *SP30B* to a temperature gradient and different exposure times.** (**a** to **e**) Activity profiles of *SP30A*, *SP30B*, *hsp30*$_{1.5kb}$ and *hsp30*$_{0.5kb}$ promoters after a short (**a** to **b** or long (**c** and **e**) heat shock treatment. Average and SD of each measurement are shown (2 to 3 biological clones, with four technical replicas for each one). A close-up of the *hsp30*$_{0.5kb}$ graph is displayed up on the right side when needed. The methodology is described in Figure S2.

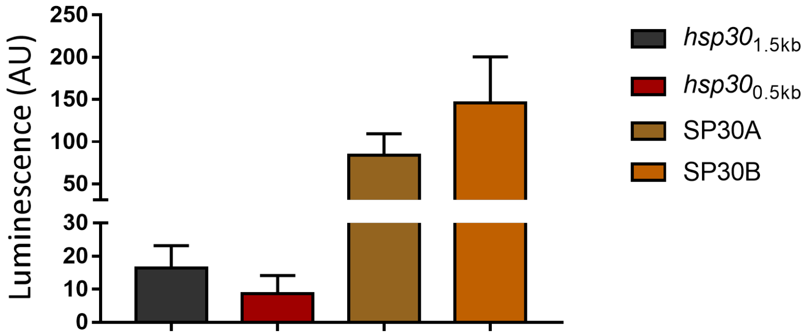


**Figure S11. Basal activity of the *SP30* promoters.** (**a**) Basal (background) luminescence levels of the *hsp30*$_{1.5kb}$, *hsp30*$_{0.5kb}$ and the *SP30* promoters. Each bar indicates the average of two or three biological clones with four independent wells each ± standard deviation (SD), and represents the behavior of two independent experiments. Values were obtained prior to delivering the heat shocks, in a 96-well plate format.

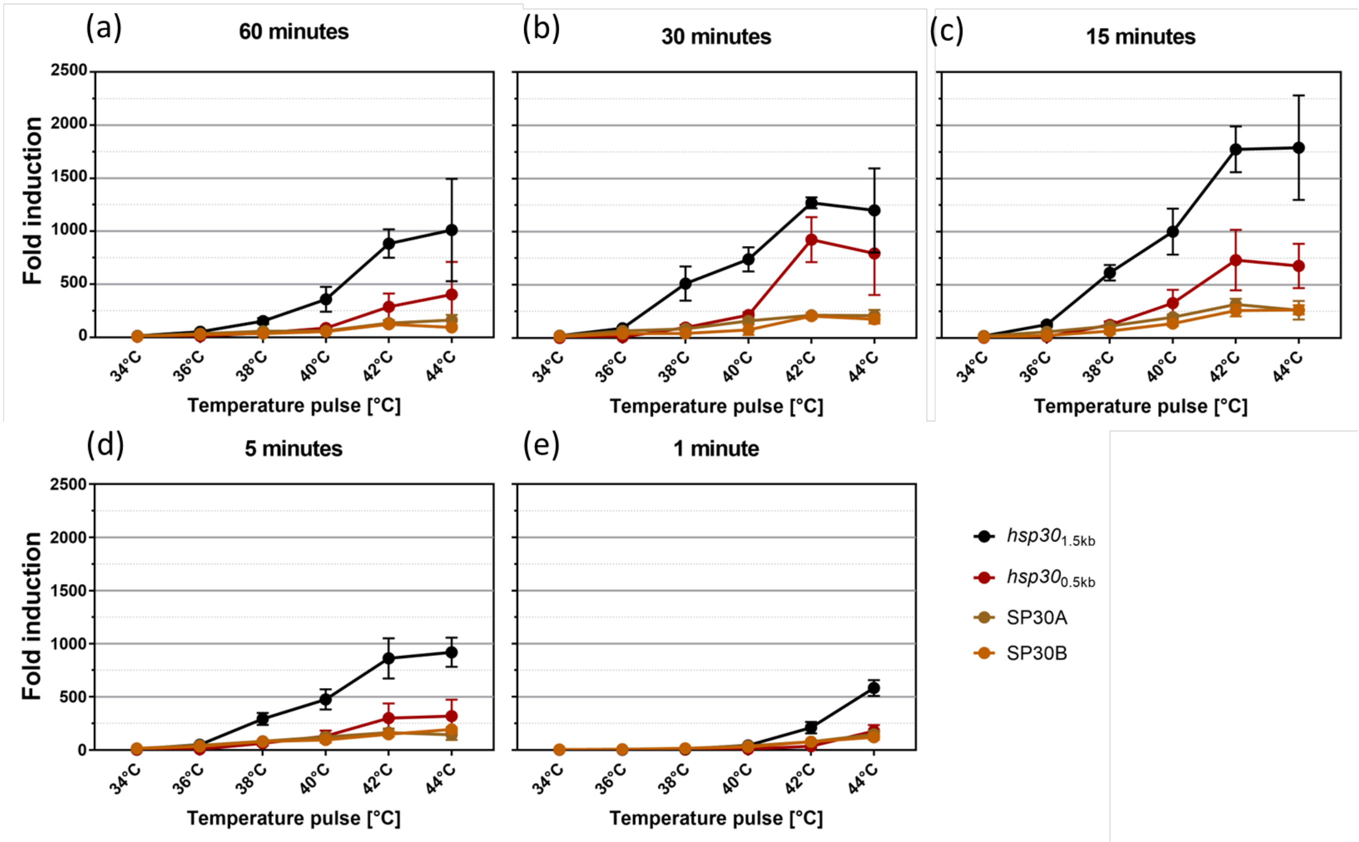


**Figure S12. Induction curves as a function of the temperature for *SP30*, *hsp30*$_{1.5kb}$ and *hsp30*$_{0.5kb}$ promoters.** (**a** to **f**) Fold induction achieved at different temperatures for *SP30A*, *SP30B*, *hsp30*$_{1.5kb}$ and *hsp30*$_{0.5kb}$ promoters during long (**a**,**b**,**c**) or short (**d**,**f**) treatments. Fold-change was calculated with the maximum luciferase expression (the average of the highest values) respect to the background levels before heat-shock treatment of each promoter region. The data was obtained from Figure S10. Each dot indicates the average of two or three biological clones with four independent wells each ± standard deviation (SD), in two independent experiments.

```
                                    HSE1
N. crassa         AGCGAATTACAGTGGCCATTCTAGAATGACCTGG-CATCTGCATCAAATGCTCCTTC-    -244
N. tetrasperma    AGTGAATTACAGTGGCCATTCTAGAATGACCTGG-CATCTGCATCAAATGCTCCTTC-    -265
N. discreta       AGTGAATTACAGTGGCTGTTCTAGAATGCCCTGG-CATTTGCGTCAAACGCTCCTTC-    -973
T. reesie         CGACAACTACAGCCGCCAGACTGGCAAGTCTTCATCATCGTCAACAACAGCCGCTGC-    -574
A. niger          CGTCTCCTGGAATGCCCGCTCTGCCGTGCGCTAA-TATCGAAACAGGGCCGAATTTC-    -1235
A. nidulans       CCAGTTTAGAAGTCTCTGCTACTGGCTGAGTAGG-CGTCTTCATCTACGGATATGCA-    -1361
A. fumigatus      ATTCCGGCCCGGTGGCGAATTCAGAGTGTAGCCAGAACCTACTATGGAGGAACATGTT    -1077
                            ...  *   .   .  .*        ..   .

                                 HSE2
N. crassa         ---GTGCAGCGCAGTGTGGAACTTTCTAGGCTGCTCCCGAAGCGTGCGATT--    -415
N. tetrasperma    ---GTGCAGTGCAGTGTGGAACTTTCTAGGCTGCTCCCGAAGCGTGCGATT--    -436
N. discreta       ---GTGCCGTGCAGTGTGGAACTTTCTAGGCTGCTCCCGAAGCGTGCGCAC--    -1147[A]
T. reesie         -GAAAGCGATG--CGGTGGAACCTTCTAGTGC-CTGCCAGAACAACCGCGC--    -1100
A. niger          AAAACCCATTG-TGAGCAATTACCTC-GCCGCGCTGTCAAAGTTTTGATTC--    -1436[A]
A. nidulans       -TACCATCTTG-ACGGCGAACTCC---CTGGTATTCCTCAGAGCAATATCGAC    -669
A. fumigatus      --GCCGCAGCG-AGCGCGAATACCACTCTGGCACTCCCGAGCAGGCCACCC--    -257
                        .   *    * . .     .       .* .. .

                              HSE3
N. crassa         -------TTTACCTC-GGCGTGAAATT-ACCAAAAAAAAAA    -567
N. tetrasperma    -------TCTACCTC-GGCGTGAAATTTACCAAAAAAGAAAA    -559
N. discreta       -------GTTACCGT-CGCACGAAATT-TCCAGAAAGTAAAC    -1251
T. reesie         GGCGAGTTCTCGCGC-AGCTTCCACTTCCCCACG--------    -105
A. niger          ----AAACCCATTGTGAGCAATTACCTCGCCGCGCTG-----    -1451[A]
A. nidulans       ---------GGGTGAAAGCAGCGAAGTTGCAAAGACCGGCGA    -523
A. fumigatus      --------GTCACGTGACCATAGAATC-ACCACGAATCATCG    -862
                                  .*    *  .  *..

                          HSE4
N. crassa         ---ATCTAGTAAACTAGCAGTAC-CAAGCTATCA    -704
N. tetrasperma    ---ATCCAGTAAACTAGCAGTAC-CAAGCTATCA    -696
N. discreta       GCCACCACTGCTACCGGCGCCAG-TAACCAG---    -251
T. reesie         ---AGCAGCAGCACCAGCAGCAGGTGCCCCAGTG    -523
A. niger          ---ATGGCGAATTCTAGAAGTCCTCTGTTCAGT-    -764
A. nidulans       ---CACGGTCCATCCAGAAGTAT-CGCCCTATAT    -1060
A. fumigatus      ---AAATCTTTTTTCAGCAATAC-CAACTCGTTA    -567
                     .          . .* .  .

                                    HSE5
N. crassa         -TACATTGAT-CACCTGGAACCTTCTTGAATGC---TCTTCAGATCTTA---    -896
N. tetrasperma    -TACATTTAT-CACCTGGAACCTTCTTGAATGC---TCTTTAGATCTTA---    -782
N. discreta       GTGC-CGTGC-AGTGTGGAACTTTCTAGGCTGC---TCCCGAAGCGTGC---    -11651[A]
T. reesie         TTAC-AGTGT-TATTTTGAGTTTCTTTTCATGT---TTTTTATTTTCCAG--    -1236
A. niger          TCGCTATCGC--TGCCAGCATCTGCCAGAACGCTC-GGCTCACTTCT-----    -816
A. nidulans       --GCTGGTGCTCAGCCAGAACCTGCTTGCCTGGAAATGTTCATGCTTTTCCA    -1305
A. fumigatus      -TGT-GATAGAATCATGGGAGATTCTCGAATCATGGCTCTGGGGTATATA--    -721
                     .             * .   * .. .  ..      . .    .
```

**Fig S13. Sequence alignment of putative *hse* sequences and surrounding regions in selected fungal *hsp30* orthologs.** "*" and "." indicate the positions conserved in 100% and 80% of sequences considered in the alignment, respectively. The bold letters indicate nucleotides that are identical to *N. crassa hse*, and the gray background shows the zone putative positions of the heat shock consensus element (HSE, 5'$TCCn_nGAAn_nTCC$3'). [A] same sequence, [B] overlapping sequences.

**Table S1**. Primers used to generate the *hsp* promoter reporters.

| Plasmid description | Primer | Sequence (5'-3') | Fragment | Backbone |
|---|---|---|---|---|
| | | **Luciferase transcritional reporters at *csr-1* locus:** | | |
| *Phsp80$_{1.1kb}$* - *luc* | oL5051 | GTCTTGTCGGGATCAGATGGACATT GAGGTaactctgatctccaaatcta | *hsp80*$_{1.1kb}$ | pLL209 |
| | oL5052 | GCCCTTCTTGATGTTCTTGGCGTCC TCCATggtGGTGgttgggcagattt | | |
| *Phsp80*$_{0.6kb}$ - *luc* | oL5053 | GTCTTGTCGGGATCAGATGGACATT GAGGTtgcggacctccagccaatca | *hsp80*$_{0.6kb}$ | pLL209 |
| | oL5052 | GCCCTTCTTGATGTTCTTGGCGTCC TCCATggtGGTGgttgggcagattt | | |
| *Phsp30*$_{1.5kb}$ - *luc* | oL5054 | GTCTTGTCGGGATCAGATGGACATT GAGGTtaaattgctgggtaatggtg | *hsp30*$_{1.5kb}$ | pLL209 |
| | oL5055 | TTGATGTTCTTGGCGTCCTCCATggt GGTGttttgactttaggtgtgtta | | |
| *Phsp30*$_{1kb}$ - *luc* | oL5056 | GTCTTGTCGGGATCAGATGGACATT GAGGTgcacgcaactgagcaaaact | *hsp30*$_{1kb}$ | pLL209 |
| | oL5055 | TTGATGTTCTTGGCGTCCTCCATggt GGTGttttgactttaggtgtgtta | | |
| *Phsp30*$_{0.5kb}$ - *luc* | oL5057 | GTCTTGTCGGGATCAGATGGACATT GAGGTcatcgagtttttcttttaaa | *hsp30*$_{0.5kb}$ | pLL209 |
| | oL5055 | TTGATGTTCTTGGCGTCCTCCATggt GGTGttttgactttaggtgtgtta | | |
| *Phsp70*$_{1.7kb}$ - *luc* | oL5058 | GTCTTGTCGGGATCAGATGGACATT GAGGTgccaaaccaatatacaaatc | *hsp70*$_{1.7kb}$ | pLL209 |
| | oL5059 | TTGATGTTCTTGGCGTCCTCCATggt GGTGtgtgaatgtgtgagatgtgc | | |
| *Phsp70*$_{1.2kb}$ - *luc* | oL5060 | GTCTTGTCGGGATCAGATGGACATT GAGGTtgcagatcaaattaccaagc | *hsp70*$_{1.2kb}$ | pLL209 |
| | oL5059 | TTGATGTTCTTGGCGTCCTCCATggt GGTGtgtgaatgtgtgagatgtgc | | |
| *Phsp70*$_{0.6kb}$ - *luc* | oL5061 | GTCTTGTCGGGATCAGATGGACATT GAGGTgcgactgatgggggaattac | *hsp70*$_{0.6kb}$ | pLL209 |
| | oL5059 | TTGATGTTCTTGGCGTCCTCCATggt GGTGtgtgaatgtgtgagatgtgc | | |
| *Pvvd*$_{3.5kb}$ - *luc* | oL3918 | GTCTTGTCGGGATCAGATGGACATT GAGGTagtggcatcaaacacaagcc | *Pvvd*$_{3.5kb}$ | pLL209 |
| | oL3919 | TTGATGTTCTTGGCGTCCTCCATggtG GTGggtgctggttatgagacagt | | |
| *Pqa-2*$_{0.6kb}$ - *luc* | AGF45 | GTCTTGTCGGGATCAGATGGACATT GAGGTtaaaacgttcgcccatcaac | *Pqa-2*$_{0.6kb}$ | pLL209 |
| | AGF46 | TTGATGTTCTTGGCGTCCTCCATggtG GTGtgtgtttggtacctctggtt | | |
| *Psp30A* - *luc* | oL6146 | GTCTTGTCGGGATCAGATGGACATT GAGGTaatacattgatcacctggaa | *SP30A* | pLL209 |
| | oL5055 | TTGATGTTCTTGGCGTCCTCCATggtG GTGttttgactttaggtgtgtta | | |

| | | | | |
|---|---|---|---|---|
| *Psp30B - luc* | oL6146 | GTCTTGTCGGGATCAGATGGACATT GAGGTcaaaacgggattcaatacat | *SP30B* | pLL209 |
| | oL5055 | TTGATGTTCTTGGCGTCCTCCATggtG GTGttttgactttaggtgtgtta | | |
| ***Phsp30*$_{0.5kb}$ controls *clr-2* transcription (locus endogenous recombination):** | | | | |
| *Phsp30*$_{0.5kb}$::Pclr-2 | oL6188 | aGCGGATAACAATTTCACACAGGAAAC AGCTCTCGGTTCTGTGACCCTCG | 5' flank | pRS426 |
| | oL6189 | acatgtaatgCATAGTACCGAGAAACTAGT CGTCCAGAAGTGTGCTCATG | | |
| | oL4543 | actagtttctcggtactatgcattacatg | *bar* cassette | |
| | LF42 | tctagactcgacagaagatg | | |
| | oL6190 | ccttcaatatCATCTTCTGTCGAGTCTAGAgc acgcaactgagcaaaact | *hsp30*$_{0.5kb}$ | |
| | oL6191 | TTGTGTAGGTATCGACATTGATGGTGC CATttttgactttaggtgtgtta | | |
| | oL2209 | ATGGCACCATCAATGTCGAT | 3' flank | |
| | oL6192 | gGTAACGCCAGGGTTTTCCCAGTCACG AGTCTTCCATGGCCTCAAGGAGC | | |

**Table S2.** *hse* identification in the *hsp30* promoter based on the *Saccharomyces cerevisiae* HSF-1 motif. The database of YeTFaSCo (http://yetfasco.ccbr.utoronto.ca/index.php) was used, where the motif ID 615 was found. To identify new *hse* or confirm the previously described on the *hsp30* promoter sequence was used FIMO of MEME suite (Version 5.4.1). A p-value < 0.001 was utilized.

| ***hse* that matched** | **Sequence (5′-3′)** | **Strand** | **p-value** |
|---|---|---|---|
| *hse5** | $^{-926}$ ttcaagaaggttcc $^{-912}$ | - | 4.2e-06 |
| | $^{-920}$ ttcttgaatgctct $^{-906}$ | + | 1.88e-05 |
| | $^{-930}$ acctggaaccttct $^{-916}$ | + | 0.000782 |
| - | $^{-881}$ tttcagggggttcg $^{-867}$ | - | 0.000883 |
| - | $^{-799}$ tttcagggggttcg $^{-785}$ | + | 0.000883 |
| - | $^{-759}$ ttatgggaagttca $^{-745}$ | + | 0.000974 |
| *hse2* | $^{-453}$ ttctaggctgctcc $^{-439}$ | + | 6.42e-05 |
| | $^{-459}$ gcctagaaagttcc $^{-445}$ | - | 0.000256 |
| *hse1* | $^{-282}$ ttctagaatgacct $^{-268}$ | + | 6.86e-05 |
| | $^{-278}$ tgccaggtcattct $^{-264}$ | - | 0.000301 |
| | $^{-288}$ ttctagaatggcca $^{-274}$ | - | 0.000968 |

*Not previously identified

**Table S3**. *hse* identification in the *hsp30* promoter based on the HSF-1 motif of *N. crassa*. The CIS-BP database (http://cisbp.ccbr.utoronto.ca/) with motif ID T243453 (HSF-1, NCU08512. To identify new *hse* or confirm the previously described sites on the *hsp30* promoter sequence using the tool "Scan single sequences for TF binding" (Motif model: PWMs-LogOdds). The default threshold was used (score equal or above 8).

| ***hse* that matched** | **Sequence (5′-3′)** | **Score** |
|---|---|---|
| *hse5** | $^{-932}$ tcacctggaaccttcttgaa $^{-912}$ | 16.94 |
| - | $^{-1238}$ gtggaagg $^{-1230}$ | 8.148 |
| - | $^{-760}$ gttatgggaagttca $^{-745}$ | 8.603 |
| - | $^{-760}$ ttagagaa $^{-745}$ | 8.862 |
| - | $^{-636}$ ggaatagtct $^{-626}$ | 8.790 |
| - | $^{-584}$ aaaaaaaaaagaa $^{-571}$ | 8.321 |
| *hse2* | $^{-463}$ gtgtggaactttctaggctg $^{-443}$ | 16.001 |
| - | $^{-316}$ cccccaagagaactccagcg $^{-296}$ | 8.338 |
| *hse1* | $^{-282}$ ttctagaa $^{-274}$ | 14.355 |
| - | $^{-240}$ ttcttcgtccttc $^{-227}$ | 11.392 |

*Not previously identified

**Table S4.** Analysis of the response obtained by inducible promoters in *N. crassa* after the corresponding stimuli

(-) Indicates that these values could not be calculated as QA remains in the media

| | Promoter | Background luminiscense (a.u.) | Max luminiscense (a.u.) | Time max luminiscense (h) | Min luminiscense post-pulse (a.u.) | Time min post-pulse (h) | Recovery time (h) | Recovery (%) | Fold induction |
|---|---|---|---|---|---|---|---|---|---|
| **Black tubes** | *vvd* | 11.3 ± 2.9 | 1694.3 ± 424.9 | 0.2 ± 0.0 | 59.3 ± 27.5 | 4.2 ± 0.2 | 4.2 ± 0.2 | 97.2 ± 1.3 | 149.9 ± 10.2 |
| | *qa* | 40.7 ± 6.2 | 1725.7 ± 470.5 | 10.8 ± 1.7 | 1286.0 ± 385.6 | - | - | - | 42.1 ± 8.2 |
| | *hsp* | 6.1 ± 3.7 | 5208.7 ± 3254.5 | 0.2 ± 0.0 | 70.3 ± 49.7 | 7.0 ± 0.0 | 7.0 ± 0.0 | 98.9 ± 0.3 | 852.0 ± 26.9 |
| **PCR tubes** | *vvd* | 87.8 ± 22.1 | 4368.7 ± 862.6 | 0.3 ± 0.2 | 326.3 ± 127.6 | 4.4 ± 1.0 | 4.3 ± 0.1 | 94.4 | 52.6 ± 18.4 |
| | *qa* | 13.2 ± 3.8 | 3267.0 ± 982.6 | 11.3 ± 0.8 | 1455.3 ± 377.0 | - | - | - | 274.6 ± 143.8 |
| | *hsp* | 4.3 ± 0.7 | 7817.7 ± 2198.0 | 2.2 ± 0.3 | 329.0 ± 13.7 | 7.0 ± 0.0 | 5.0 ± 0.3 | 95.6 ± 1.2 | 1804.8 ± 459.7 |
